# Improved Supra-Organization Is A Conformational And Functional Adaptation Of Respiratory Complexes

**DOI:** 10.1101/2020.01.03.892737

**Authors:** Shivali Rawat, Suparna Ghosh, Debodyuti Mondal, Valpadashi Anusha, Swasti Raychaudhuri

## Abstract

Multiple surveillance mechanisms accelerate proteasome mediated degradation of misfolded proteins to prevent protein aggregation inside and outside mitochondria. But how cells safeguard mitochondrial function despite increased protein aggregation during proteasome inactivation? Here, using two-dimensional complexome profiling, we extensively characterize the dynamic states of respiratory complexes (RCs) in proteasome-inhibited cells. We report that RC-subunits are increasingly integrated into supra-organizations to optimize catalytic activity simultaneous to their aggregation inside mitochondria. Complex-II (CII) and CV are incorporated into oligomers. CI, CIII, and CIV subcomplexes are associated into holocomplexes followed by integration into supercomplexes. Time-course experiments reveal that the core (CI+CIII_2_) stoichiometry of supercomplex (I+III_2_+IV) is preserved during early-stress while CIV composition varies. Simultaneously, increased CI-activity suggests conformational optimization of supercomplexes for better function. Re-establishment of steady-state stoichiometry and relative increase in supercomplex-quantity consolidates functional adaptation during prolonged proteasome-inhibition. Together, we name this pre-emptive adaptive mechanism as ‘improved Supra-organization of Respiratory Complexes’ (iSRC). We find that iSRC is active in multiple protein-unfolding stresses, in multiple cell-types that differ in proteostatic and metabolic demands, and reversible upon stress-withdrawal.

## INTRODUCTION

Cellular adaptation to adverse external conditions is implemented at multiple levels. Stress-responsive expression of chaperones is widely observed but not a rapid adaptation due to delays associated with transcription and translation (Zhang et al, 2015). Instead, optimization of existing proteins represents a faster mechanism to fight stress during initial impact and serves as activation-signal for transcriptional stress responses. Conformational alteration, allosteric or non-allosteric regulation, phase separation and reversible aggregation, posttranslational modifications, translocation etc. include dynamic mechanisms to perceive stress, to trigger stress response pathways and to reorganize cellular components to withstand stress (Franzmann & Alberti, 2019; Gsponer & Babu, 2012; Iida et al, 1986; Ohn & Anderson, 2010; Winter & Jakob, 2004).

Another immediate cellular response to any kind of stress is to shut-down energy-expensive metabolic processes. For example, protein synthesis is highly energy-expensive and is globally reduced in response to stress (Lane & Martin, 2010). Nevertheless, spending energy in cap-independent translation of stress-chaperones, in degradation of misfolded proteins by ATP-dependent ubiquitin-proteasome system (UPS) or in refolding of unfolded proteins by chaperones remain indispensable. Therefore, preserving optimal activities of energy-production machineries and investing available energy in adaptive mechanisms is essential to survive stress and to mount defence responses.

Oxidative phosphorylation is the most efficient energy generation pathway in cell. Four respiratory complexes (RCs), NADH-ubiquinone oxidoreductase (Complex I), succinate-ubiquinone oxidoreductase (Complex II), ubiquinone-cytochrome-c oxidoreductase (Complex III), and cytochrome-c-oxidase (Complex IV) generate an electrochemical gradient across inner mitochondrial membrane to convert ADP to ATP by ATP synthase (Complex V) (Mitchell & Moyle, 1968). RCs are composed of proteins of dual genetic origin and assemble in mitochondrial matrix. Nuclear-encoded RC-subunits associate with mitochondria-encoded subunits; first into subcomplexes and then into holocomplexes (Fernandez-Vizarra & Zeviani, 2015; Guerrero-Castillo et al, 2017; Signes & Fernandez-Vizarra, 2018; Vidoni et al, 2017). Cellular protein quality control capacity prevents accumulation of unassembled subunits and controls fidelity of RC-assembly by selectively preserving founding sub-complexes for on-demand assembly. We have found that unassembled RC-subunits are aggregation-prone proteins(Rawat et al, 2019). Interestingly, despite aggregation, oxygen consumption by individual RCs remained preserved during short-term proteasome inhibition.

RCs are functionally independent in terms of substrate utilization. However, large proportion of CI, CIII and CIV co-operate as interacting enzymes to facilitate efficient substrate oxidation. Such supra-organizations of RCs are known as ‘respiratory supercomplexes’ or ‘respirasomes’ (Gu et al, 2016; Guo et al, 2018). CII can also be part of a respiratory megacomplex but predominantly present as tetramer while CV exists as dimeric and oligomeric complexes along cristae edges (Bezawork-Geleta et al, 2018; Guo et al, 2018; Wittig & Schagger, 2009). Association of individual RCs into supercomplexes is a dynamic event and depends on the metabolic status of the cell (Porras & Bai, 2015).

Studies on plasticity of RCs is so far restricted to blue-native PAGE followed by western blotting that reports distribution of 1-2 subunits per complex. Furthermore, switch of RC-subunits from subcomplexes to holocomplex and supercomplexes is not resolved in these experiments (Balsa et al, 2019; Greggio et al, 2017). Here, we established an iBAQ and SILAC-based two-dimensional complexome profiling protocol for comprehensive subunit-wise mapping of relative distribution of subunits between different RC-organizations in multiple conditions. Using this strategy, we find that formation of high molecular weight supercomplexes is favoured over free individual complexes and subcomplexes in response to accumulation of protein-aggregates inside mitochondria during proteasome-inactivation. Our results suggest that this dynamic posttranslational defence mechanism; hereafter referred to as improved Supra-organization of Respiratory Complexes (iSRC), represents a reversible spatiotemporal proteome adaptation driven by conformational and functional optimization of RC-subunits.

## RESULTS

### DISTRIBUTION OF RC-SUBUNITS IN RESPIRATORY COMPLEXES

#### RC-subunits in mitochondrial fraction during proteasome inhibition

We have recently reported that Respiratory Complex (RC) subunits represent a group of highly expressed chaperone-sensitive proteins that form reversible aggregates in Neuro2a cells during short-term proteasome inhibition (8 hr). Interestingly, despite aggregation, mitochondrial import of RC-subunits was not blocked (Rawat et al, 2019). Continuously imported RC-subunits may have three fates inside mitochondria during proteasome-inhibition. They may remain as soluble individual subunits and as part of RCs; precipitate into insoluble fraction; or be degraded by other mitochondrial proteases or mitophagy. To investigate, we performed SILAC-based quantification of mitochondrial fractions of Neuro2a cells after proteasome-inhibitor MG132 treatment (5 µM). Abundance of RC-subunits remained unaltered in total and soluble fraction after short-term (8 hr) treatment (**Figure S1A and Table S1**). However, several CI, CII and two CIII-subunits were increased in insoluble fraction suggesting accumulation of aggregation-prone subunits. Exceptionally, CIV-subunits were not increased rather reduced in insoluble fraction. Depletion of many RC-subunits in soluble fraction at 24 hr was prominent by increasing blue intensity in heatmap (**Figure S1A**). Simultaneously, abundance of multiple RC-subunits was reduced in total mitochondrial fraction probably due to disposal of unstable subunits by mitochondrial proteases (Ruan et al, 2017) or active autophagy in Neuro2a cells during prolonged proteasome-inhibition (Rawat et al, 2019).

#### Distribution of RC-subunits in different RC-organizations

Oxygen consumption by individual RCs was only marginally dropped after 8 hr MG132 (Rawat et al, 2019) even though multiple RC-subunits accumulated in the insoluble fraction (**Figure S1A**). Therefore, we performed native-PAGE followed by western blotting to correlate structural integrity with reducing solubility and consequent reorganization of RCs inside mitochondria. Probing with CI-subunit Ndufs1 revealed presence of CI in multiple species with different masses (**Figure 1A**). The band at ∼800 kDa indicated the CI-holocomplex (CI-HC) while the upper bands represented CI-supercomplexes (CI-SCs) (**Figure1A**) (Guerrero-Castillo et al, 2017; Jha et al, 2016). We also observed increase in SCs at 8 hr MG132 while CI-HC remained similar to control. On the other hand, slight drop in intensity for all CI-bands was noticed at 24 hr. Ndufs1-level was not increased in total and soluble mitochondrial fractions (**Figure S1A**) suggesting its increase in CI-SCs at 8 hr could be due to redistribution from lower molecular weight subcomplexes (SubCs).

**Figure 1.**
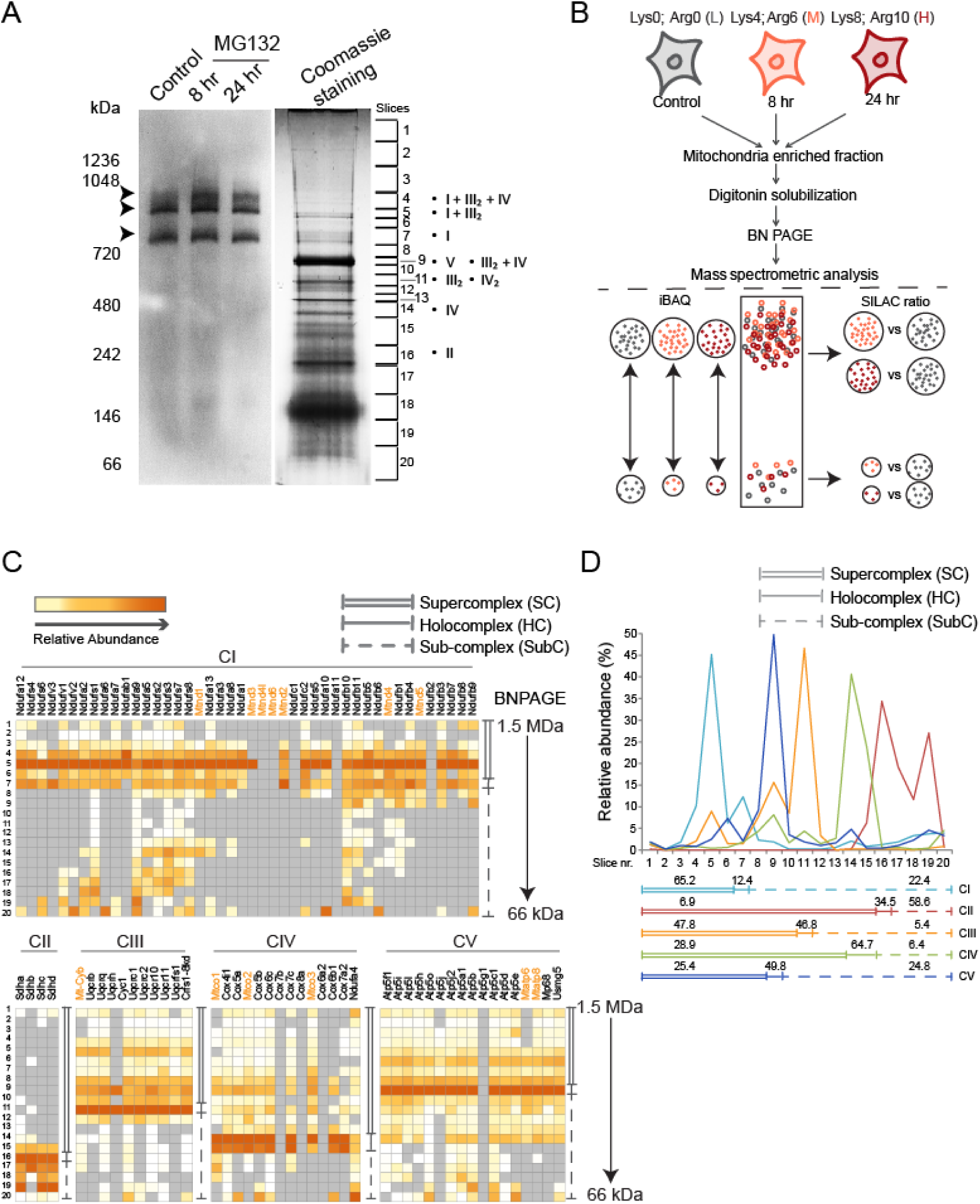
Distribution of RC-subunits in different RC-organizations. **A) Left:** Digitonin soluble mitochondrial fraction was loaded on native-PAGE, blotted with Ndufs1-antibody and imaged. Arrows indicate CI-bands. **Right:** Similar gel was stained with coomassie brilliant blue showing separation of mitochondrial protein complexes. The gel slices processed for mass spectrometry are marked and numbered. Dots indicate RCs identified in respective slices. **B)** Flowchart describing iBAQ and SILAC-based two-dimensional complexome profiling strategy using BN-PAGE. Each dot represents one RC-subunit. Cells are treated with MG132 (5 μM) for indicated time-points. Color-code for SILAC-labelled peptides. **C)** Heatmap showing relative abundance of RC-subunits as determined by iBAQ values in unstressed Neuro2a cells at different slices of BN-PAGE. Mitochondria-encoded subunits are shown in orange. n=2. **D)** Relative abundance distribution profile of RCs across the BN-PAGE. The abundance of all identified subunits for respective RCs in a given slice was summed together. Relative abundance in that slice was estimated in comparison to cumulative abundance of all subunits of the same RC across all slices and distribution profile was plotted. The plots represent subunit distribution in three sub-groups: supercomplex (SC), holocomplex (HC) and subcomplexes (SubC). Percentage abundance in each group is indicated. n =2.

Ndufs1 in subcomplexes (below CI-HC at ∼800kDa) was not detected in western-blots (**Figure 1A**). Therefore, we improvised an iBAQ and SILAC-based complexome profiling protocol to comprehensively map the distribution of RC-subunits in SCs, HCs and SubCs. The Blue Native-PAGE (BN-PAGE) was divided in 20 slices according to coomassie-staining keeping the Ndufs1-bands separate, and quantitative mass spectrometry was performed (**Figure 1A**). In this strategy, SILAC-ratios report increase or decrease of abundance of RC-subunits within the same slice of BN-PAGE due to MG132-treatment. Simultaneously, relative iBAQ intensities of isotopic peptides across the slices provide distribution profiles of SCs, HCs and SubCs in multiple conditions (**Figure 1B**).

First, we calculated slice-wise absolute abundance of RC-subunits in unstressed Neuro2a cells as per iBAQ algorithm (Schwanhausser et al, 2011). Consistent with in-gel CI activity, slice 7 (∼800kDa) contained CI subunits with moderate iBAQ-values suggesting presence of fully assembled CI-HC (**Figure 1C and Table S2A**). CI interacts with the CIII dimer (CIII_2_) to form SC (I+III_2_), or with both CIII_2_ and CIV to constitute bigger SCs (I+III_2_+IV), (I+III_2_+IV_2_) etc. (Jha et al, 2016; Milenkovic et al, 2017). Bright CI activity band and higher cumulative intensity of CI and CIII-subunits in slice 5 indicated that majority of CI in Neuro2a cells was associated with CIII (**Figure 1C, Table S2A-B**). We also found CI-activity bands in slice 4 (∼1000 kDa). CI, CIII and CIV subunits were identified here suggesting presence of SC (I+III_2_+IV) but in less abundance than SC (I+III_2_) (**Figure 1C, Table S2A-B**). CI, CIII and CIV subunits were either absent or present in low intensities in top three slices suggesting absence of further high molecular weight SCs in Neuro2a cells (**Figure 1C**).

CV subunits were most abundant in slice 9 (∼ 700 kDa) corresponding to CV-HC but also in higher slices (slice 6 and 8) suggesting multimeric organizations (**Figure 1C and Table S2B**). Abundance of both CIII and CIV in slice 9 suggested presence of SC (III_2_+IV) (**Figure 1C**). CIII and CIV subunits were also identified in slice 11 representing dimeric HCs (Jha et al, 2016; Lapuente-Brun et al, 2013). Monomeric CIV-HC was found in slice 14-15 while CII-HC was in slice 16 (**Figure 1C**). Slice 9-15 contained multiple Q and P_D_ module components indicating presence of intermediate CI-SubCs (Guerrero-Castillo et al, 2017). Core subunits of N module of CI (Ndufv1, Ndufv2 and Ndufs1) were found in slice 16-20 (∼ 150-250 kDa) along with several Q and P_D_ module components suggesting multiple small sub-complexes (**Figure 1C**). Sub-complexes of CIII and CV were also present between slice 13-20 as evident by abundance of their constituting subunits (**Figure 1C**) (Fernandez-Vizarra & Zeviani, 2015; Vidoni et al, 2017). Taken together, relative distribution of iBAQ intensities across the BN-PAGE suggested that most of the CI-subunits were present within SCs while CIII and CIV subunits were also abundant in HCs (**Figure 1D**). CII-and CV subunits were distributed between HC and SubCs although 25.4% of CV-subunits were found to be part of multimeric organizations.

### IMPROVED ASSEMBLY OF RESPIRATORY COMPLEXES DURING PROTEASOME INHIBITION

#### SILAC-based quantification of RC-subunits in MG132 treated Neuro2a cells

Multiple N and Q module SubCs of CI below slice 9 of the BN-PAGE was found to be reduced with MG132 treatment (**Figure 2A**). Small SubCs of P_D_ module below slice 16 were also reduced whereas intermediate SubCs (between slice 9-16) were comparatively increased at 8 hr (**Figure 2A and Table S2C**). Uqcrc1, Uqcrc2 and Cyc1 together form an CIII-SubC (Fernandez-Vizarra & Zeviani, 2015). Both Uqcrc1 and Uqcrc2 were reduced in slice 19-20. CIV SubCs composed of (Cox4i1+Cox5a) and (Cox5b+Cox6c) (Signes & Fernandez-Vizarra, 2018) were depleted in slice 19-20. Stator stalk SubC of CV composed of Atp5f1, Atp5l and Atp5i (Fujikawa et al, 2015) in slice 17-18 was also reduced by proteasome inhibition. Complex II is constituted through independent maturation of Sdha, Sdhb and (Sdhc + Sdhd) (Signes & Fernandez-Vizarra, 2018). We found Sdha, Sdhc and Sdhd in slice 19 to be reduced by MG132 treatment (**Figure 2A and Table S2D**).

**Figure 2.**
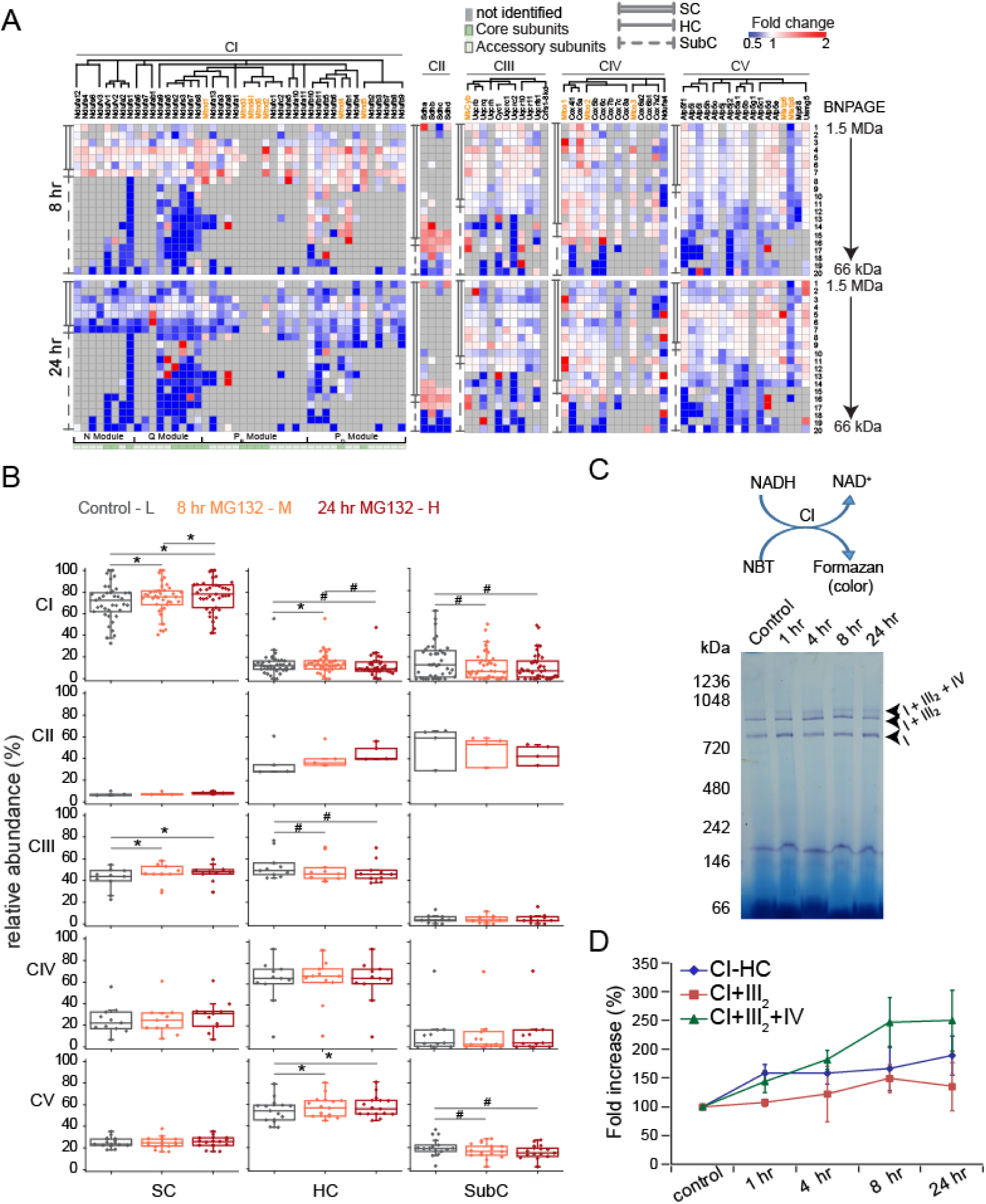
Improved assembly of RCs during proteasome inhibition in Neuro2a cells. **A)** Heatmap showing SILAC based fold changes in abundance of RC-subunits in MG132 (5 µM) treated Neuro2A cells at different slices of BN PAGE. The subunits are clustered as per the assembly pathways described earlier (Fernandez-Vizarra & Zeviani, 2015; Fujikawa et al, 2015; Guerrero-Castillo et al, 2017; Vidoni et al, 2017). Mitochondria-encoded subunits are shown in orange. n =2. **B)** Neuro2a cells were treated with 5 µM MG132 for 8 and 24 hr. Box-plots showing relative abundance (iBAQ) distribution of individual RC-subunits across three sub-groups. iBAQ value of a subunit in a given slice (n) was divided by summation of iBAQ values of the same subunit in all 20 slices. Each dot represents individual subunits of respective RCs. % relative abundance for SCs, HCs and SubCs are according to the sub-groups shown in **Figure 1D**. * or # indicates increase or decrease respectively, where p< 0.05 one-way ANOVA repeated measures, Bonferroni’s post hoc test. SILAC isotopic peptides representing different experimental conditions are colour coded; L - light, M- medium, H- heavy. n =2. **C)** In-gel activity of NADH dehydrogenase (CI). **Top**: Experimental scheme. **Bottom**: Cells were treated with 5 µM MG132 for indicated time-points, Digitonin soluble mitochondrial fraction was prepared and loaded on native-PAGE, incubated with substrates and imaged. Arrows indicate CI activity-bands. **D)** Fold increase in activity of NADH dehydrogenase (CI) at increasing time-points of 5 µM MG132. Activity band intensities were calculated and plotted. Error bars indicate SD from three independent experiments.

SILAC-ratios for all CII subunits were increased in the slice 15-16 at 8 hr MG132 suggesting consolidation of CII-HC (**Figure 2A**). Similarly, CI subunits were either increased or remained unchanged at slice 7 and above where CI-HC and SCs were found. Most of the CI and CIII subunits composing SC (I+III_2_) at slice 5 slightly increased or remained unchanged at 8 hr. CI, CIII and CIV subunits were prominently increased at slice 4 at 8 hr suggesting improvement of SC (I+III_2_+IV) (**Figure 2A**).

In contrast to short-term proteasome inhibition (8 hr), abundance of CI-HC and CI-SCs was reduced at 24 hr MG132 (**Figure 2A**). Subunits in CIII and CV-HCs at slice 11 and 9 remained unchanged or marginally reduced. CII and few CIV subunits displayed increased abundance at slice 15-17 and 14-15, respectively in their HCs. Majority of the CIII and CIV subunits remained comparatively unchanged at slice 9 suggesting protection for SC (III_2_+IV) (**Figure 2A**).

#### iBAQ-based relative distribution of RC-subunits in MG132 treated Neuro2a cells

Significant increase in relative distribution of CI subunits into SCs with increasing time of MG132 was confirmed as per iBAQ intensity profile of isotopic peptides (**Figure 2B and S2A**). CI-HC was found to be slightly improved at 8 hr but dropped with time. On the other hand, drop in CI-subunits in SubCs was prominent with MG132. Increase of SCs and drop in SubCs was most prominent for N and Q modules (**Figure S2B**) that protrude out from inner mitochondrial membrane (IMM) towards the hydrophilic environment of matrix (Letts et al, 2016). Membrane-resident P module was comparatively unchanged.

Increased incorporation of CII and CV-subunits into HCs due to proteasome inhibition was observed with concomitant drop of their SubCs (**Figure 2B and S2A**). On the other hand, relative abundance of CIII-HC was significantly reduced and with simultaneous improvement of SC (**Figure 2B**). Association of CIII and CIV-subunits into SC (III_2_+IV) in slice 9 was increasingly stronger with MG132 treatment (**Figure S2A**). Incorporation of CI, CIII and CIV-subunits into SCs in slice 4-5 was also improved. Thus, despite overall depletion of RC-subunits at longer proteasome inhibition (24 hr) as depicted by SILAC heatmap; iBAQ data suggested favoured distribution of remaining soluble subunits towards SC-formation at the expense of subassemblies. Hereafter, we term this phenomenon as improved Supra-organization of Respiratory Complexes (iSRC).

#### iSRC - a functional adaptation

In order to investigate whether improved physical association of RC-subunits into super-quaternary structures was a functional adaption in stressed cells, we performed in-gel CI activity assay (**Figure 2C**). CI-HC activity-band became brighter than control already by 1 hr MG132 suggesting better substrate utilization with stress (**Figure 2C-D**). SC (I+III_2_) was more abundant than SC (I+III_2_+IV) (**Figure 1D**) but its activity was not increased except 8 hr (**Figure 2C-D**). Intensity of SC (I+III_2_+IV) activity band was gradually brighter with increasing time of stress. Maximum activity was achieved by 8 hr when increased integration of RC-subunits into SCs was also observed (**Figure 2C-D**). Activity-band intensities for CI-HC and SC (I+III_2_+IV) remained brighter than control unstressed cells even at 24 hr MG132 (**Figure 2C-D**). This suggested that despite depletion of RC-subunits across all RC-organizations (**Figure 2A**); CI-HC and SCs remained optimized for better function during prolonged proteasome inhibition.

### iSRC - A MULTI-STEP PROTEOME ADAPTATION

#### Time kinetics of iSRC

Many stress-adaptive proteome reorganization mechanisms are instantaneous. For example, small stress granules emerge in cytoplasm within 3-4 min stress and then progressively fuse with each other to form bigger granules (Nadezhdina et al, 2010). Moreover, improved CI-activity at SC (I+III_2_+IV) indicated a possibility of increased incorporation of RC-subunits into SCs by already 1 hr MG132. Therefore, to investigate kinetics of iSRC we performed complexome profiling experiments at 1 and 4 hr MG132 in Neuro2a cells. In spite of variations in BN-PAGE slicing by visually inspecting coomassie stained bands, relative distribution profile of RC-subunits in SCs, HCs and SubCs remained reproducible in control unstressed cells between experiments **(Figure S2A and S3A)**. Surprisingly, unlike 8 hr MG132 (**Figure 2B)**, improvement of SCs was not apparent at 1 and 4 hr. Instead, distribution of RC-subunits into CIII and CV-SCs was reduced (**Figure 3A, S3A and Table S3**). CI-HC was reduced with a concomitant increase in SubCs (**Figure 3A**). Distribution of CI-subunits was marginally increased in Q- and P_D_ module SubCs (**Figure S3B**).

**Figure 3.**
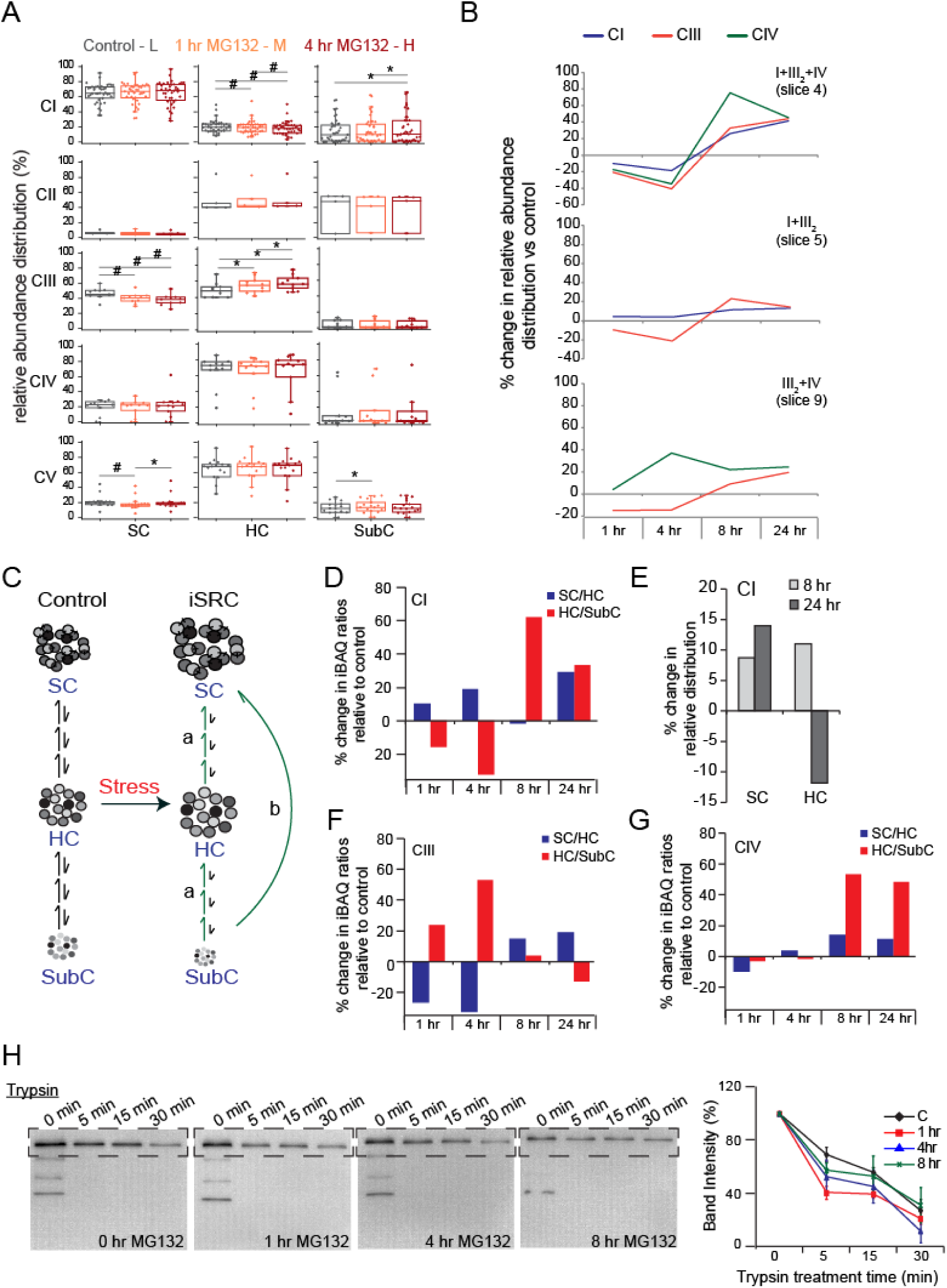
Multi-step mechanism of iSRC. **A)** Neuro2a cells were treated with 5 µM MG132 for 1 and 4 hr. Box-plots showing relative abundance (iBAQ) distribution of individual RC-subunits of all RCs across three sub-groups. Each dot represents individual subunit of respective RC. * or # indicates increase or decrease respectively, where p< 0.05 one-way ANOVA repeated measures, Bonferroni’s post hoc test. Experimental conditions and SILAC isotopic peptides are colour coded in Figure. n=2 **B)** Percent change in relative abundance for RCs within SCs in different slices of BN-PAGE at indicated time-points of MG132 compared to control unstressed cells. **C)** Model depicting possible mechanisms of SubC integration into SCs during iSRC. SubCs may first integrate into HCs and in a second step HCs may integrate into SCs (route a). Or, in SubCs may directly integrate into SCs (route b). Or, one-step and multi-step mechanisms may work simultaneously. **D)** Percent change iBAQ ratio of CI SC/HC and CI HC/SubC in Neuro2a cells treated with 5 µM MG132 for indicated time-points compared to control unstressed cells. **E)** Percent change in relative abundance distribution in CI-SC and CI-HC in Neuro2a cells treated with 5 µM MG132 for indicated time-points compared to control unstressed cells. **F-G)** Percent change iBAQ ratio of SC/HC and HC/SubC of CIII and CIV in Neuro2a cells treated with 5 µM MG132 for indicated time-points compared to control unstressed cells. **H)** Limited proteolysis of Ndufs1. Neuro2a cells were treated with 5 µM MG132 for 0, 1, 4 and 8 hr. Digitonin-solublised mitochondrial fraction (2 μg) from treated cells was digested with 100 ng Trypsin for indicated time-points and loaded on SDS-PAGE. Western blot was performed using anti-Ndufs1 antibody. Band intensities from the boxed-region of the blots were calculated and plotted (right). Error bars indicate SD from three independent experiments.

Similar to iBAQ data, increasing blue intensity in slice 4-7 of SILAC heatmap suggested abundance of CI-HC and SCs was reduced during early-stress especially at 4 hr (**Figure S3C**). SubCs remained unaltered or increased below slice 13-14 at 1 hr MG132. However, P_D_ module SubCs in slice 14-19 and Q-module SubCs at slice 19-20 were increased in abundance at 4 hr (**Figure S3C**). CIII-HC at slice 11-12 was also increased although CIII-SCs and SubCs were reduced in abundance at 4 hr (**Figure S3C**). Overall drop in abundance of CIV subunits was observed throughout SILAC-heatmap (**Figure S3B**).

Further, time-kinetics data revealed disproportionate distribution of individual RCs within different SCs at early time-points of MG132 (**Figure 3B**). SC (I+III_2_+IV) in slice 4 was most dynamic. CI and III increased proportionately in slice 4 at 8 hr but CIV fold-increase was ∼ 30% excess than CI and CIII (**Figure 3B**). This suggested although a large population of SCs remained at (I+III_2_+IV) stoichiometry, a sub-population of SCs with increased CIV units may also had emerged at this stage. Similarly, disproportionate CIV-distribution compared to CIII at 1 and 4 hr was observed at slice 9 (**Figure 3B**). However, changes in relative distribution for CI, III and IV absolutely overlapped in slice 4, 5 and 9 at 24 hr suggesting that despite overall depletion (**Figure 2A**) of CI, III and IV-subunits relatively increased and consolidated at SC (I+III_2_+IV), (I+III_2_) and (III_2_+IV) as per steady-state stoichiometry at this time-point (**Figure 3B**).

#### A stepwise integration of SubC>HC>SC

It is proposed that during biogenesis CI, CIII and CIV-subunits are first incorporated into HCs and then integrated into SCs (route a; **Figure 3C**) (Guerrero-Castillo et al, 2017). This mechanism may remain consistent for iSRC. Alternatively, SubCs may directly integrate into SCs (route b, **Figure 3C**). Only 12.4% of CI-subunits were found within HC in Neuro2a cells suggesting it to be a transient CI-assembly that rapidly integrated into SCs or dissociated into SubCs (**Figure 1D**). CI-HC was reduced at 1 hr with a concomitant increase in SubCs resulting in decreased CI HC/SubC ratio (**Figure 3A and D**). CI SC/HC ratio was increased simultaneously as relative distribution of CI-subunits in SCs remained unchanged. Further drop in CI-HC was observed at 4 hr (**Figure 3A**). Simultaneously, CI-SubCs increased further and HC/SubC ratio decreased sharply (**Figure 3D**). CI-SC and HC both were proportionately increased at 8 hr (**Figure 3E**). CI SC/HC ratio remained unchanged; however, increased HC/SubC ratio suggested incorporation of SubCs into HC (**Figure 3D**). Integration of CI-HC into SCs increased at 24 hr with a concomitant drop in CI-HC (**Figure 3E**). Therefore, CI SC/HC ratio was increased at 24 hr MG132 (**Figure 3D**).

In contrast to CI, only 5% of CIII subunits were found in SubCs (**Figure 1D**). Increase in CIII HC/SubC iBAQ ratio at 1 and 4 hr MG132 suggested increased incorporation of subunits into HC (**Figure 3A and F**). Furthermore, drop in CIII SC/HC ratio indicated simultaneous disintegration of SCs. CIII SC/HC was increased at 8 hr and HC/SubC remained close to control (**Figure 3F**). On the other hand, unchanged SC/HC and HC/SubC ratios suggested that relative distribution of CIV-subunits between SubC, HC and SCs remained similar to control at both 1 and 4 hr (**Figure 3G**). Both SC/HC and HC/SubC ratios were increased for CIV at 8 hr indicating synchronized incorporation of subunits into higher assemblies. Thus, iSRC also followed two-step SubC>HC>SC mechanism similar to biogenesis; however, each complex showed distinct integration kinetics.

To probe the dynamics of RC-organizations further, we performed limited proteolysis of mitochondria-enriched fractions from MG132 treated cells and blotted for N-module subunit Ndufs1. Abundance of Ndufs1 was found to be increased in mitochondrial insoluble fraction by 8 hr MG132 confirming its instability with stress (**Figure S1A**). As plotted in **Figure 3H**, Ndufs1 was also more accessible to protease-digestion in MG132-treated fractions compared to control unstressed cells. Rapid digestion of Ndufs1 was observed after 1 hr MG132 as evident by drop in band intensity after 5 min protease treatment suggesting maximum conformational perturbation at that time-point. Thus, together the time-kinetics experiments revealed that iSRC was not an instantaneous proteome-adaptation rather RC-subunits underwent extensive reorganization before their improved incorporation into SCs.

### iSRC SUBSIDES DURING STRESS RECOVERY

Generally, stress-adaptive proteome reorganizations are reversible. Stress granules disappear within hours of stress-recovery (Kedersha & Anderson, 2007). Protein aggregates also resolubilize after withdrawal of heat-stress (Wallace et al, 2015). Aggregates of over-expressed CI-subunit Ndufa5 emerged at 8 hr MG132 and disappeared after MG132-withdrawal (**Figure 4A**). Intensity of CI-activity bands was also decreased by 16 hr MG132 withdrawal; but not up to pre-stress condition (**Figure 4B-C**). Particularly, SC (I+III_2_+IV) remained at an improved-functional state.

**Figure 4.**
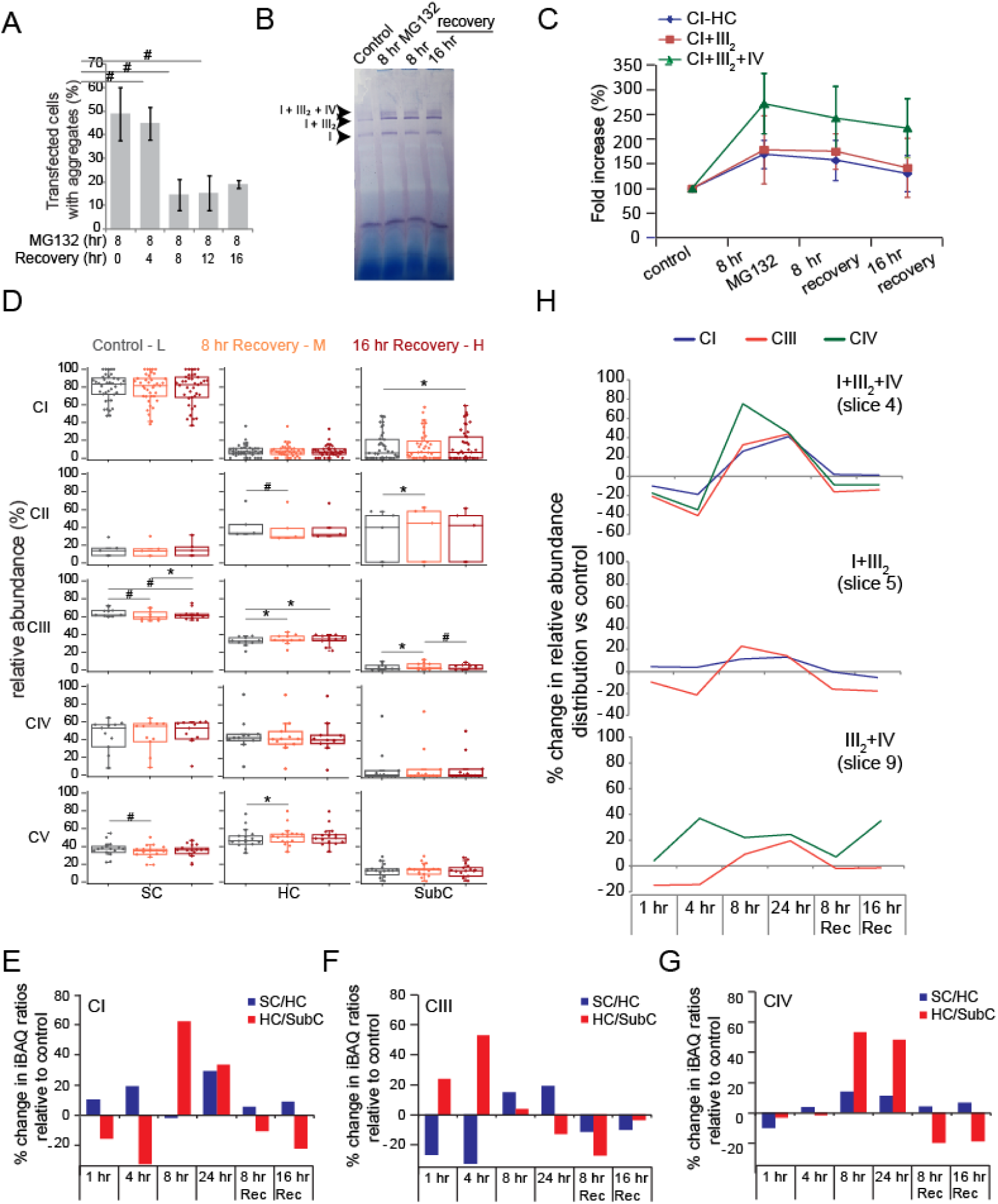
Reversion of iSRC. **A)** Graph showing percentage of Neuro2a cells with Ndufa5-EGFP aggregates. Cells were transfected with Ndufa5-EGFP followed by 5 µM MG132 treatment for 8 hr. MG132 was withdrawn by adding fresh media and cell counting was done at indicated time-points. Error bars indicate SD; 10 different fields with approximately 25 transfected cells have been counted. # indicates p< 0.05 in Student’s t-test. **B)** In-gel activity of NADH dehydrogenase (CI). Cells were stressed for 8 hr using 5 µM MG132 followed by recovery for indicated time-points. Arrows indicate activity bands. **C)** Fold increase in activity of NADH dehydrogenase (CI) at indicated time-points of 5 µM MG132 treatment. Activity band intensities were calculated and plotted. Error bars indicate SD from three independent experiments. **D)** Box-plots showing relative abundance (iBAQ) distribution of RC-subunits across three sub-groups. Each dot represents individual subunit of respective RC. * or # indicates increase or decrease respectively, where p< 0.05 one-way ANOVA repeated measures, Bonferroni’s post hoc test. Experimental conditions and SILAC isotopic peptides are colour coded in Figure. **E-G)** Percent change iBAQ ratio of SC/HC and HC/SubC of CI, CIII and CIV, respectively in Neuro2a cells treated with 5 µM MG132 for indicated time-points compared to control unstressed cells. **H)** Percent change in relative abundance for RCs within SCs in BN-PAGE slices at different time-points of MG132 compared to control unstressed cells.

Decrease in SubCs and simultaneous increase in HCs and SCs was a signature of iSRC at 8 hr MG132 (**Figure 2**). This trend was reversed during recovery. As per iBAQ calculation of isotopic peptides, distribution of subunits in CI, CII and CIV-SCs was restored to normal equilibrium whereas CIII-SCs and multimeric organizations of CV were reduced compared to control unstressed cells (**Figure 4D, S4A, Table S4A-B**). Increase in CII-HC level and decrease in CIII-HC observed during iSRC (**Figure 2**) was reversed after MG132-withdrawal (**Figure 4A and S4A**). SILAC heatmap also confirmed comparatively more SubCs than high molecular weight assemblies during recovery (**Figure S4B, Table S4C-D**). Abundance of subunits in CIII-HC (slice 11-12) was increased while SCs were reduced (**Figure S4B**). CIII-SubCs were also improved during early recovery. As depicted by increasing blue intensity of the heatmap, higher order RCs were more reduced at 16 hr recovery compared to 8 hr (**Figure S4B**). Improved catalytic activity of SC (I+III_2_+IV) at 16 hr (**Figure 4C**) suggested continued presence of functionally optimized SCs despite reduction in quantity (slice 4; **Figure S4B**).

Dissociation of SCs was also found during early-stress (1 and 4 hr MG132); however, we found subtle differences with stress-recovery. CI-Q module was marginally redistributed between HC and SubCs at early-stress (**Figure S3B**). Instead, CI-N-module was increased while P_D_ module was prominently decreased in SCs during recovery (**Figure S4C**). Also, CI-HC was reduced during early-stress (**Figure 3A**). In contrast, CI-HC restored equilibrium during recovery (**Figure 4D**). However, CI SC/HC and HC/SubC iBAQ ratios remained similar during early-stress and recovery (**Figure 4E**). CIII SC/HC and HC/SubC ratios reached close to equilibrium at 16 hr recovery (**Figure 4F**). On the other hand, distribution of CIV subunits between SubCs, HC and SCs maintained equilibrium at early-stress while HC/SubC ratio decreased during recovery (**Figure 4D-G**). Further, relative changes in distribution for individual RCs suggested restoration of equilibrium during recovery although CIII was marginally under-proportion at SC (I+III_2_+IV) and (I+III_2_) (**Figure 4H**). Disproportionate relative increase in CIV at slice 9 suggested extensive re-organization of SC (III_2_+IV). Thus, mechanism of iSRC was distinct for individual RCs in terms of kinetics of integration into SCs during stress and dissociation during recovery.

### STABILIZATION OF SCs AS GENERAL ADAPTIVE STRATEGY DURING PROTEOSTASIS-STRESS

#### iSRC during heat stress

Increased protease-sensitivity of Ndufs1at early-stage of proteasome inhibition tempted us to investigate iSRC in other protein-destabilizing stresses. Active degradation machinery continuously degrades unfolded proteins in heat-stressed cells (Raychaudhuri et al, 2014). Therefore, drop in Ndufs1 level in total and soluble mitochondrial fraction of heat stressed Neuro2a cells (42°C for 2 hr) could be due to degradation of unfolded Ndufs1 (**Figure 5A**). In agreement, we also observed depletion of subunits at CI-HC and SCs in SILAC heatmap of complexome profiling (**Figure S5A**). Ndufs1 protein load was restored in mitochondrial fraction by 2 hr recovery (37°C for 2 hr after heat stress; **Figure 5A**) possibly with the assistance of increasing heat-shock chaperones. Similarly, decreasing blue intensity in SCs and HCs suggested restoration of RC-subunits during recovery (**Figure S5A**). CI-activity within SCs was also protected in heat-stressed cells (**Figure 5B**).

**Figure 5.**
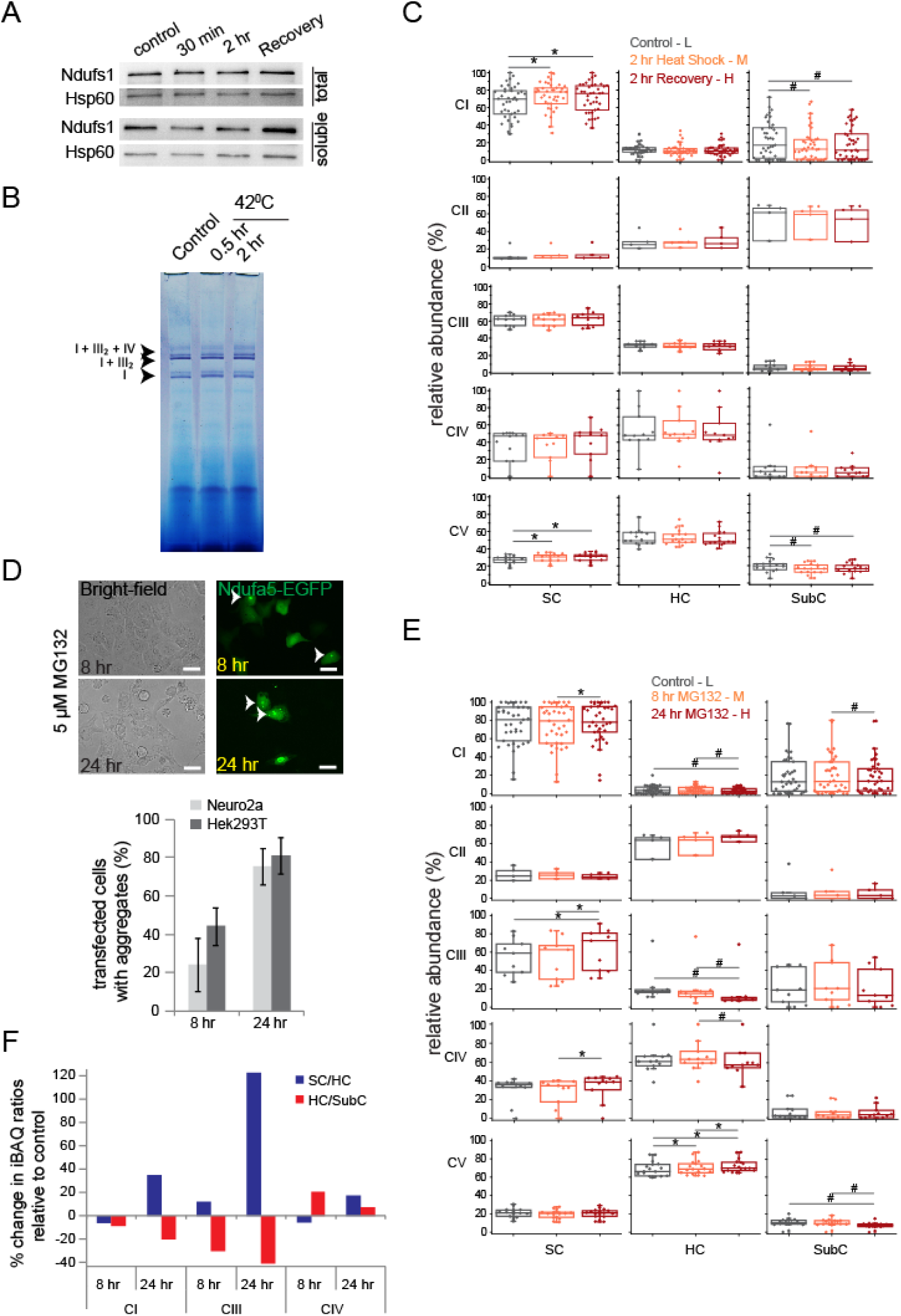
iSRC as general adaptive strategy. **A)** Neuro2a cells were heat stressed (42°C) for 30 min and 2 hr followed by recovery at 37°C for 2 hr. Total and soluble mitochondrial fraction from treated cells was loaded on SDS-PAGE and western blot was performed using anti-Ndufs1 antibody. Hsp60 served as loading control. **B)** In-gel activity of NADH dehydrogenase (CI). Cells were heat stressed (42°C) for 30 min and 2 hr. Arrows indicate activity bands. **C)** Box-plots showing relative abundance (iBAQ) distribution of individual RC-subunits across three sub-groups. Neuro2a cells were heat stressed (42°C for 2 hr) followed by recovery (37°C for 2 hr). Each dot represents individual subunit of respective RC. * or # indicates increase or decrease respectively, where p< 0.05 one-way ANOVA repeated measures, Bonferroni’s post hoc test. Experimental conditions and SILAC isotopic peptides are colour coded in Figure. **D) Top:** microscopy images of 5 µM MG132 treated Ndufa5-EGFP transfected Hek293T cells. After 24 hr of transfection, cells were treated with MG132 for indicated time-points. Arrows indicate aggregates. Scale-bar - 50 µm. **Bottom:** percentage of Neuro2a and Hek293T cells with aggregates of Ndufa5-EGFP after 5 µM MG132 treatment. Error bars indicate SD; 10 different fields with approximately 25 transfected cells have been counted. **E)** Box-plots showing relative abundance (iBAQ) distribution of individual RC-subunits of all RCs across the three sub-groups. Hek293T cells were treated with 5 µM MG132 for 8 and 24 hr. Each dot represents individual subunit of respective RC. * or # indicates increase or decrease respectively, where p< 0.05 one-way ANOVA repeated measures, Bonferroni’s post hoc test. Isotopic peptides representing different experimental conditions colour coded in Figure. **F)** Percent change iBAQ ratio of SC/HC and HC/SubC of CI, CIII and CIV in Hek293T cells treated with 5 µM MG132 for indicated time-points compared to control unstressed cells.

iBAQ redistribution of RC-subunits was indicative of stabilization of SCs. Increase in CI-SCs and oligomeric organization of CV was observed with a concomitant drop in their SubCs (**Figure 5C, S5B, Table S5A-B**). Similar to MG132 stress, N- and Q-modules within CI-SCs was prominently increased during heat-stress (**Figure S5C**). Strikingly, we did not observe simultaneous increase in CIII and CIV suggesting a non-stoichiometric distribution of RCs within SCs during heat stress. Thus, iSRC in heat stressed cells was limited to CI and CV but protected SC-function despite overall depletion of subunits (**Figure S5A**).

#### iSRC in Hek293T cells

Finally, we investigated iSRC in Hek293T cells. SC-organizations vary in different cell-lines depending on source-organisms or tissues, differential expression of RC-subunits and assembly factors, proteostasis capacity inside and outside mitochondria, etc. For example, association of CIII_2_ with CIV into SCs is regulated by longer isoform of Cox7A2l or supercomplex assembly factor 1 (SCAF1) (Lapuente-Brun et al, 2013). We have identified only the shorter isoform of Cox7A2l in Neuro2a cells (**Figure S6A**) and large SCs including (I+III_2_+IV_2_), (I+III_2_+IV_3_) and (III_2_+IV_2_) were absent (**Figure 1C**). Although Cox7A2l - knockout human cells also lack SC (III_2_+IV); it was present in Neuro2a (**Figure 1C-D**). In contrast, high molecular weight SCs including respiratory megacomplex (I_2_+III_2_+IV_2_) are reported to be present in Human embryonic kidney cell line Hek293T (Guo et al, 2018). Despite discrepancy in SC-organization, primary amino acid sequences of RC-subunits are nearly-identical in mouse and human cells (Fearnley & Walker, 1992) and similarly aggregation-prone. For example, mouse Ndufa5 is ∼83% sequence-conserved with human Ndufa5 and found to form aggregates upon proteasome inhibition when over-expressed in Hek293T cells (**Figure 5D and S6B**).

The distribution profile of RCs in Hek293T cells was different compared to Neuro2a (**Figure 1D and S6C**). CI and CIII subunits were majorly present in high molecular weight SCs including the megacomplex (I_2_+III_2_+IV_2_) in slice 2-4. In comparison to Neuro2a, abundance of CI and CIII subunits in HCs (slice 7 and 11, respectively) and in SC (I+III_2_) in slice 5 was less. CIV level was more in the top four slices although CIV-HC remained highly abundant like Neuro2a cells in slice 13-14. Similarly, CII-HC and CV-HC were abundant in slice 16 and 9, respectively (**Figure S6C, Table S6A-B**).

iSRC remained consistent in Hek293T cells. Increasing red intensity in top slices of SILAC heatmap suggested comparatively more stable and consolidated high molecular SCs in Hek293T cells, even after 24 hr MG132 (**Figure S6D**). The megacomplex (I_2_+III_2_+IV_2_) was markedly improved in Slice 2. Multiple intermediate CI-subcomplexes also remained stabilized in slice 8, 9 and 10. CII-HC remained unchanged in slice 16 and SubCs were reduced in lower slices (**Figure S6D, Table S6C-D**). CIII-SubCs in slice 12-20 were improved at 8 hr, however, were depleted at 24 hr when most of the CIII subunits accumulated in SCs in slice 2-3 (**Figure S6D**). CIV-HC and SCs were found to be improved in slice 14 and above; more prominently at 24 hr (**Figure S6D**). Improvement of higher order assemblies of CV was not apparent in SILAC heatmap. Rather, CV subunits remained unchanged at 8 hr or slightly reduced throughout the BN-PAGE. CV-SubCs below slice 9 were highly reduced at 24 hr MG132 treatment although higher assemblies were comparatively protected (**Figure S6D**).

Relative isotopic iBAQ distribution of RC-subunits suggested that CI-SCs, particularly the megacomplex (I_2_+III_2_+IV_2_) in slice 2, were significantly improved (**Figure 5E, S6C, Table S6A-B**). Simultaneously, CI-SubCs were reduced with time. Similar to Neuro2a cells, increase of SCs and drop in SubCs was prominent for N- and Q-module subunits at 24 hr (**Figure S6E**). CII- and CV-HCs were improved in slice 16 and 9, respectively. Simultaneously, presence of CV subunits in SubCs was significantly reduced with time (**Figure 5E**). CIII and CIV-SCs were increased at slice 2-3, predominantly at 24 hr at the expense of HCs (**Figure 5E**). Increase in SC/HC iBAQ ratios for CI, CIII and CIV was not observed at 8 hr but only after 24 hr MG132 in Hek293T cells indicating slower iSRC kinetics compared to Neuro2a (**Figure 5F**).

## DISCUSSION

Cells evolved multiple surveillance strategies including mPOS, UPRam, mitoCPR etc. to prevent accumulation of protein-aggregates inside and outside mitochondria (Boos et al, 2019; Wang & Chen, 2015; Weidberg & Amon, 2018; Wrobel et al, 2015). These mechanisms work via transcriptional upregulation of protein quality control machinery that eventually aid to degradation of misfolded proteins. In absence of efficient degradation, RC-subunits are the first group of proteins to accumulate and form aggregates (Rawat et al, 2019). In this work, we demonstrate aggregation of RC-subunits in mitochondrial fraction of proteasome-inhibited cells. Further, we identify and characterize a posttranslational response referred to as iSRC that triggers integration of remaining soluble RC-subunits inside mitochondria into supercomplexes (SCs). Relative increase in SCs persists during extreme proteome-destabilizing conditions of prolonged proteasome inhibition or acute heat stress. As a result, catalytic function of RCs is protected and improved. Our data confirms that the equilibrium shift of RC-subunits from smaller assemblies to super-quaternary structures is a subtle but consistent, organized and reversible adaptation.

### iSRC - CONFORMATIONAL AND FUNCTIONAL OPTIMIZATION OF RCs

Respiratory complexes are multimeric enzymes. Increased substrate utilization by enzymes could be a direct outcome of increasing enzyme concentration or due to conformational optimization for faster substrate-consumption. As per limited proteolysis, RC-subunits enjoy conformational flexibility at early stages of proteasome inhibition. Complexome profiling data indicates simultaneous dissociation of a fraction of ‘loosely associated’ RC-subunits from SCs and HCs. However, rather than perturbing function, partial disintegration ensures optimum conformation of SCs and HCs to achieve catalytic advantage.

Time kinetics by complexome profiling also suggests dynamic stoichiometry of SCs at early-stress. SC (I+III_2_+IV) in slice 4 of BN-PAGE represents the largest SC in Neuro2a cells. At 8 hr proteasome inhibition, excess CIV-subunits in this slice argue for a transient SC-subpopulation in which more than one CIV-unit is associated with (I+III_2_). CIV is the most dynamic RC in terms of SC-constitution. Cryo-electron microscopy revealed two distinct architectures for SC (I+III_2_+IV); in the ‘tight’ structure CIV is attached to both CI and CIII. ‘Loose’ structure refers to contact between CI and CIV only (Letts et al, 2016). Two separate CIV monomers associated with (I+III_2_) in situ has also been reported in mammalian cells (Davies et al, 2017). Further, in-gel CI-activity assays revealed (I+III_2_) to be associated with up to 3 CIV-units in multiple studies (Jha et al, 2016). Interestingly, we do not always observe a distinct activity band for SC (I+III_2_+IV_2_) or higher in our in-gel assays but a smeary appearance was noticed. As SILAC-based quantification protocol eliminates biochemical artefacts of SC-preparation from differently stressed samples; disproportionate increase of CIV in slice 4 suggests a dynamic population of SC (I+III_2_+IV_1or2_) that is not well-resolved in our BN-PAGEs. Similarly, excess CIV in slice 9 at early-stress suggests a subpopulation of (III_2_+IV_2_). Thus, in contrast to rigid super-quaternary structures with fixed stoichiometry; we observe plasticity and inequivalence of constituent RCs suggesting ‘quinary’ SC-organizations at early-stress (Gierasch & Gershenson, 2009; Hingorani & Gierasch, 2014; McConkey, 1982). These dynamic SC-ensembles interconvert at multiple stages of assembly or disassembly to re-establish steady-state stoichiometry of SC (I+III_2_+IV) and (III_2_+IV) with time (**Figure 6**).

**Figure 6.**
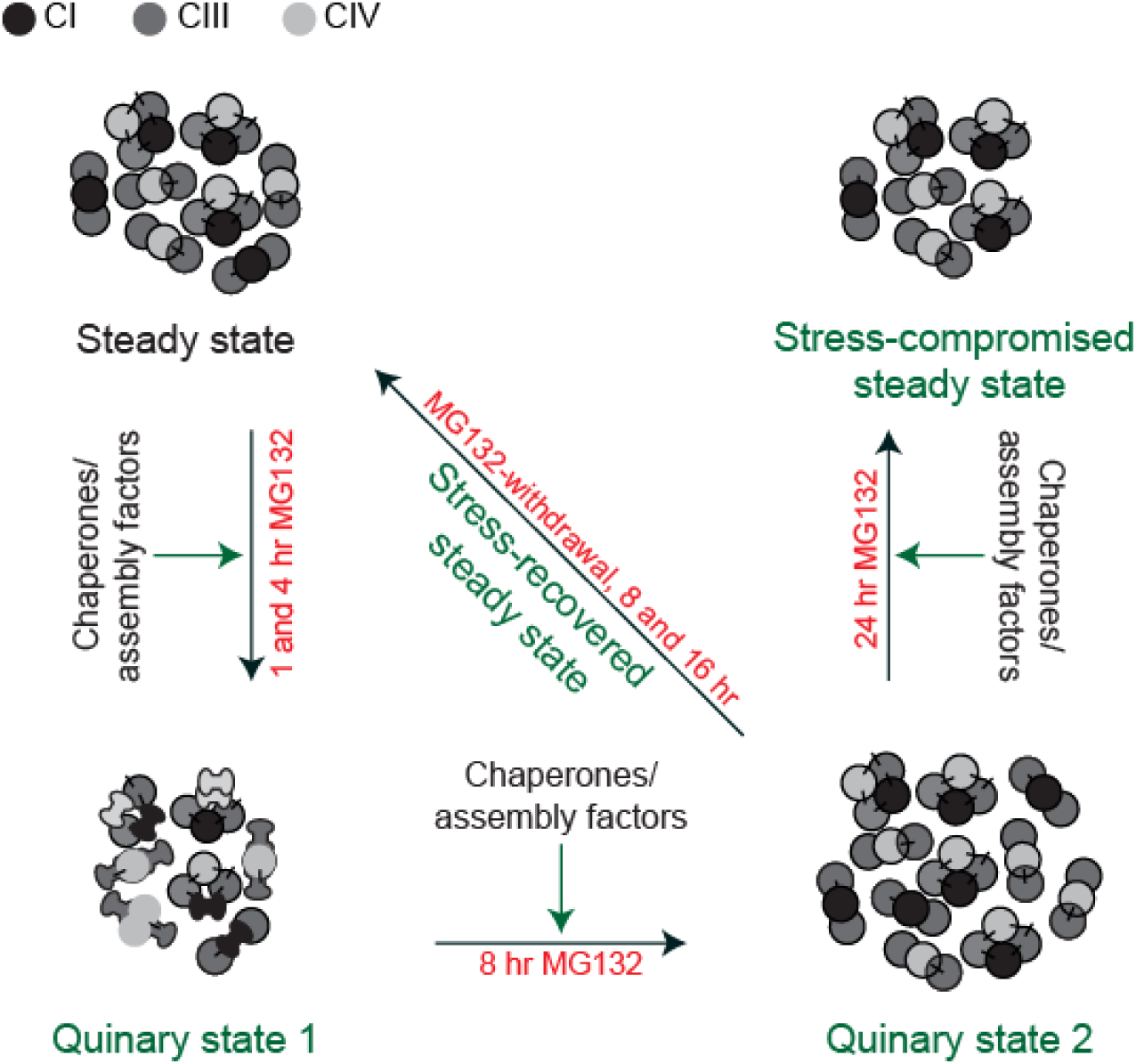
Model depicting iSRC - dynamic adaptation of respiratory supercomplexes (SCs) during proteasome inhibition as revealed by two-dimensional complexome profiling. Five distinct states of SCs were observed. Steady state: Cells grown in normal culture conditions in absence of MG132. SCs are attributed to literature described stoichiometry – SC (I+III_2_+IV) and SC (I+III_2_). Quinary state 1: Partial disintegration of SCs during early-MG132 (5 µM, 1 and 4 hr) as revealed by both iBAQ and SILAC. Excess CIV in SCs indicates non-steady state stoichiometry. Conformational flexibility as probed by limited proteolysis. Increased catalytic activity. Quinary state 2: 8 hr MG132. Increased SC-quantity as revealed by both iBAQ and SILAC. Excess CIV in SCs indicates non-steady state stoichiometry. Increased catalytic activity. Stress-compromised steady state: 24 hr MG132. Overall drop in SCs, HCs and SubCs as revealed by SILAC. Relative protection of SCs over SubCs as revealed by iBAQ. Re-establishment of steady state stoichiometry. Increased catalytic activity. Stress-recovered steady state: MG132 withdrawal. Restoration of steady state abundance of SCs as per SILAC. Steady state stoichiometry as per iBAQ. Increased catalytic activity.

CI devoid of N-module has been shown to associate with CIII to form partially assembled SC-intermediates in assembly-factor depleted cells (Calvaruso et al, 2012; Stroud et al, 2016). It is also possible that CIV_2_ directly associate with similar partially assembled SC-intermediates that move together with (I+III_2_+IV) or (III_2_+IV) in slice 4 or 9 of BN-PAGE. However, partially assembled CI-intermediates may not be abundant in Neuro2a cells as SILAC-ratios suggest that reduction of CI-subunits during early-stress is not limited to any particular module.

Our results identify early-stress and recovery periods as most dynamic states of RC-organizations. RC-subunits dissociate yet SCs and HCs remain functionally improved. However, at early-stress SCs experience dynamic stoichiometry while at recovery remain closed to steady-state stoichiometry. At these ‘quinary-state’ of SC-organizations (Powers & Balch, 2013; Roth & Balch, 2013), free RC-subunits, SubCs and RCs may interact with assembly chaperones to stabilize their soluble forms and to favour stage-specific stoichiometry. Hence, iSRC may not be only driven by proximity-based weak interactions as proposed by Blaza et al (Blaza et al, 2014) and likely to be under elaborate regulation. Recently longer isoform of Cox7A2l (SCAF1) has been described to improve SC-organization in U2OS cells (Balsa et al, 2019). This isoform is not present in Neuro2a but iSRC is active suggesting cell-specific assembly mechanisms for iSRC are yet to be deciphered. Also, in situ structural analysis at early-stress and recovery is expected to reveal novel dynamics of functional and conformational cross-talks (Letts et al, 2019).

## OUTLOOK

Endoplasmic Reticulum stress has been reported to stimulate SC-formation (Balsa et al, 2019). Proteasome inhibition protects RC-assembly factor COA7 from degradation and improves CIV function in models of mitochondrial leukoencephalopathy (Mohanraj et al, 2019). Our data also suggests that triggering iSRC by mild proteostasis stress may offer an opportunity to ameliorate OxPhos-deficiency in mitochondrial diseases. On the other hand, many protein complexes assemble co-translationally to prevent aggregation of free subunits (Kamenova et al, 2019; Shiber et al, 2018). Nuclear-encoded RC-subunits do not enjoy that opportunity as these are imported from cytoplasm before their assembly inside mitochondria. Therefore, a considerable fraction of free RC-subunits and SubCs always exist inside mitochondria with aggregation-prone interaction surfaces exposed. Controlled and specific integration of free sub-assemblies into SCs offers an indirect fitness-benefit during proteostasis-stress by preventing non-functional interactions and subsequent aggregation. The two-dimensional complexome profiling strategy described here could prove powerful to investigate spatiotemporal organization of many such dynamic protein interactions explaining myriad of biological functions.

## MATERIALS AND METHODS

### Constructs

Ndufa5 was PCR-amplified from Neuro2a cDNA and cloned into pcDNA4/TO EGFP using the restriction enzymes KpnI and XhoI, as described earlier (Rawat et al, 2019).

### Cell culture and microscopy

Neuro2a and Hek293T cells were maintained in Dulbecco’s Modified Eagle’s Medium (Gibco) supplemented with 10% fetal bovine serum (FBS) (Gibco) and 90 U/ml penicillin (Sigma) - 50 µg/ml streptomycin (Sigma) at 37°C and 5% CO_2_. Transfection of cells was performed with Lipofectamine 3000 reagent (Invitrogen) according to the manufacturer’s protocol for 24 hr. For aggregation kinetics, random fields were captured using FLoid Cell Imaging Station (Thermo Scientific) at the different time-points. Student’s t-test was performed for significance calculation.

For SILAC based mass spectrometry - cells were grown in SILAC DMEM (Thermo Scientific) supplemented with 10% dialyzed FBS (Gibco), 90 U/ml penicillin (Sigma) – 50 µg/ml streptomycin (Sigma) and either Light [L-Lysine 2HCl / L-Arginine HCl (Lys0/Arg0)] or Medium [L-Lysine 2HCl (4,4,5,5-D4) / L-Arginine HCl (^13c^_6_) (Lys4/Arg6)] or Heavy [L-Lysine 2HCl (^13c^_6_, ^15^N_2_) / L-Arginine HCl (^13c^_6_, ^15^N_4_) (Lys8/Arg10)] isotopes of lysine and arginine (Thermo Scientific).

### Mitochondrial fraction preparation

Mitochondria enriched fraction preparation from cultured cell lines was performed according to Schägger (1995) and Rebeca Acín-Pérez (2008), with some modifications (Acin-Perez et al, 2008; Schagger, 1995). Briefly, ∼10 million cells were pooled together and pellet was washed twice with PBS. Cell pellet was frozen at −80°C and homogenized in a dounce homogenizer with about 10 cell pellet volumes of homogenizing buffer, buffer A (83 mM Sucrose, 10 mM MOPS [pH 7.2]). After adding an equal volume of buffer B (250 mM Sucrose, 30 mM MOPS [pH 7.2]), nuclei and unbroken cells were removed by centrifugation at 1000g during 5 min at 4°C. Mitochondria enriched fraction was collected from the supernatant by centrifuging at 12000g during 5 min and washed twice under the same conditions with buffer C (320 mM Sucrose, 1 mM EDTA, 10 mM Tris-HCl [pH 7.4]). The pellet was resuspended in buffer D (1 M 6-aminohexanoic acid, 50 mM Bis-Tris HCl [pH 7.0]), and estimated by Pierce Coomassie Plus (Bradford) Assay Reagent (Thermo Scientific).

### Mass spectrometry sample preparation from mitochondrial fraction

Mitochondrial fractions were lysed in NP-40 lysis buffer (50 mM Tris-HCl [pH 7.8], 150 mM NaCl, 1% NP-40, 0.25% Sodium deoxycholate, 1 mM EDTA, protease inhibitor cocktail (Roche)) at 4°C for 45 min with intermittent vortexing. After centrifugation at 12000g for 15 min at 4°C, supernatant was collected as the soluble fraction. The remaining pellet was washed twice with 1X PBS and boiled in 4X SDS loading buffer (0.2 M Tris-HCl [pH 6.8], 8% SDS, 0.05 M EDTA, 4% 2-mercaptoethanol, 40% glycerol, 0.8% Bromophenol blue) for 15 min to get insoluble fraction. Total fraction was prepared by directly dissolving and boiling the mitochondria-enriched pellet in 4X SDS loading buffer for 15 min. The fractions were separated on NuPAGE 4-12% Bis-Tris Protein Gels (Invitrogen). The gel was run in MES buffer (100 mM MES, 100 mM Tris-HCl, 2 mM EDTA, 7 mM SDS) at 200V for 40 min, fixed and stained with coomassie brilliant blue. Preparation of gel slices, reduction, alkylation, and in-gel protein digestion were carried out as described by (Shevchenko et al, 1996). Finally, peptides were desalted and enriched according to Rappsilber et al (Rappsilber et al, 2003).

### BN-PAGE and In-gel activity

Mitochondrial fractions were solubilized by adding 8 g digitonin (Sigma) per gm of mitochondria and incubating 5 min in ice. The supernatant was collected after a 30 min centrifugation at 13000g. 5% coomassie brilliant blue G-250 dye in 1 M 6-aminohexanoic acid was added to 1/3 the final volume of supernatant. Glycerol was added at final concentration of 10% and sample was loaded onto NativePAGE 3-12% Bis-Tris Protein Gels (Invitrogen). Native PAGE was run using anode buffer (50 mM Bis-Tris HCl [pH 7.0]) and blue cathode buffer (50 mM Tricine, 15 mM Bis-Tris HCl [7.0], 0.02% coomassie brilliant blue G-250) at 100 V until samples entered separating gel and then at 150 V until dye-front reached 1/3 length of entire gel. At this point, blue cathode buffer was replaced with dye-free cathode buffer for remainder of run at 200 V until dye-front reached end of gel. The gel was incubated overnight at RT in freshly prepared CI NADH dehydrogenase substrate solution (5 mM Tris-HCl [pH 7.4], 0.1 mg/ml NADH and 2.5 mg/ml Nitrotetrazolium Blue chloride). Appearance of violet bands is indicative of CI activity. The reaction was stopped with 10% acetic acid, washed with water and imaged (Jha et al, 2016). Band intensity was calculated using ImageJ.

### Complexome profiling

Mitochondria were isolated from SILAC-labelled control and treated cells and pooled together. 100-120 µg digitonin-solubilized mitochondria were loaded onto NativePAGE 3-12% Bis-Tris Protein Gels (Invitrogen) and run as described above. The gel was fixed and stained with coomassie brilliant blue R-250 and cut into 20 slices for mass spectrometry. Preparation of gel slices, reduction, alkylation, and in-gel protein digestion were carried out as described by (Shevchenko et al, 1996). Finally, peptides were desalted and enriched according to (Rappsilber et al, 2003).

### LC-MS/MS

Peptides eluted after desalting were dissolved in 2% formic acid and sonicated for 5 min. Samples were analysed on Q-Exactive HF (Thermo Scientific) and were separated on a EASY-Spray PepMap RSLC C18 Column (75 μm × 15 cm; 3 μm) using a 60 min linear gradient of the mobile phase [5% ACN containing 0.2% formic acid (buffer-A) and 95% ACN containing 0.2% formic acid (buffer-B)] at a flow rate of 300 nL/min. Full scan MS spectra (from *m*/*z* 400–1650) were acquired followed by MS/MS scans of Top 10 peptides with charge states 2 or higher. The mass spectrometry based proteomics data of all these experiments have been deposited to the ProteomeXchange Consortium via the PRIDE partner repository (Vizcaino et al, 2014) with the dataset identifier < PXD014427 >.

### Peptide identification and statistical analysis

For peptide identification, raw MS data files of individual slices were analysed separately using MaxQuant (Ver. 1.3.0.5) (Cox & Mann, 2008) and searched against Swissprot database of *Mus musculus* (release 2018.10 with 25208 entries) or *Homo sapiens* (release 2019.03 with 42419 entries) and a database of known contaminants. MaxQuant used a decoy version of the specified database to adjust the false discovery rates for proteins and peptides below 1%. The search parameters included constant modification of cysteine by carbamidomethylation, enzyme specificity trypsin, multiplicity set to 3 with Lys4 and Arg6 as medium label and Lys8 and Arg10 as heavy label. Other parameters included minimum peptide for identification 1, minimum ratio count 1, re-quantify option selected. iBAQ option was selected to compute abundance of the proteins. Bioinformatics and statistical analysis was performed in Perseus environment (Ver. 1.5.2.4) (Tyanova et al, 2016).

For experiments with two biological repeats, average values generated as per MaxQuant algorithm were used for further analysis. For total and soluble fractions M/L and H/L ratios, were converted into log2 space, and mean ratios and standard deviations were calculated for each data set (Coombs et al, 2010). The log2 M/L and H/L ratio of each protein were converted into a z-score, using the following formula:

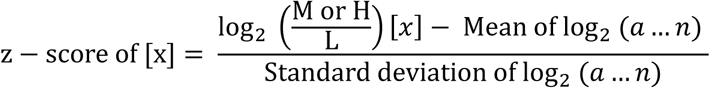

where x is a single protein in the data set population (a….n). The z-score was a measure of how many Standard deviation (σ) units, the log2 M/L or H/L ratio of the protein was away from the population mean. A z-score ≥ 1.96σ represented that differential expression of the protein lied outside the 95% confidence interval and were considered to be significant. As insoluble fraction showed skewed distribution, mean and standard deviation of total fraction was used to normalize and calculate z-score.

### Abundance and statistical analysis

Protein abundance in different gel slices was calculated by selecting iBAQ option in MaxQuant tool. Thereafter, all calculations were carried out separately for light, medium or heavy labeled peptides. Abundance of all identified subunits in a given slice (n) was summed together for respective RCs. Relative abundance was estimated in comparison to cumulative abundance of all subunits of the same RC identified across all slices and distribution profile was plotted.

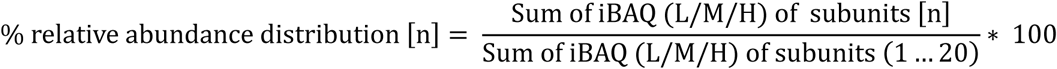

The % relative abundances were summed together to calculate distribution in three sub-groups namely Supercomplex (SC), holocomplex (HC) and subcomplex (SubC).

To calculate relative abundance distribution of individual subunits (x) across slices, iBAQ value of the subunit in a given slice (n) was divided by summation of iBAQ values of the same subunit in all 20 slices.

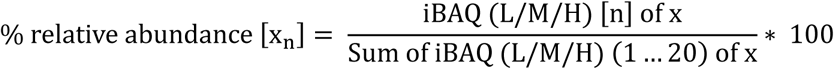

Statistical analyses were performed in OriginPro 8 (OriginLab Corporation, Northampton, MA, USA). Differences among multiple groups were compared using one-way ANOVA repeated measures, Bonferroni’s post hoc test. Student’s t-test was performed for normally distributed datasets. p < 0.05 was considered statistically significant.

Percent changes in SC/HC and HC/SubC iBAQ ratio was calculated according to the following formula:

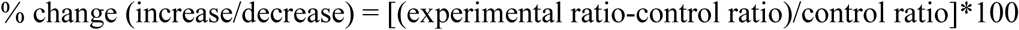

### Limited proteolysis and western blotting

Digitonin solubilised mitochondrial fraction (2 μg) from control and treated cells was digested with 100 ng Trypsin for indicated time points. Reaction was stopped by adding 4X SDS loading buffer and boiling the samples. The samples were separated by SDS-PAGE and transferred onto 0.2 μm PVDF membrane (Bio-rad) for 90 min at 300 mA using the Mini-Trans Blot cell system (Bio-rad). Membranes were probed by appropriate primary and secondary antibodies (**Supplementary Methods Table 1**) and imaged using documentation system (Vilber Lourmat). Band intensity was calculated using ImageJ.

## Supplementary Methods Table 1

**Antibody list:**

| Antibody Name | Company | Catalog number | Dilution |
| --- | --- | --- | --- |
| Rabbit monoclonal Anti-Ndufs1 antibody [EPR11521(B)] | Abcam | Cat#ab169540 | 1:1000 |
| Anti-Rabbit IgG (whole molecule) (peroxidase conjugate) | Sigma | Cat#A6154 | 1:20000 |
| Anti-Hsp60 antibody [LK-1] | Abcam | Cat#ab59457 | 1:1000 |
| Anti-Mouse IgG (whole molecule) (peroxidase conjugate) | Sigma | Cat#A4416 | 1:20000 |

