## Supplementary figures and legends for "Improved Supra-Organization Is A Conformational And Functional Adaptation Of Respiratory Complexes"

### SUPPLEMENTARY FIGURES AND FIGURE LEGENDS

Figure S1

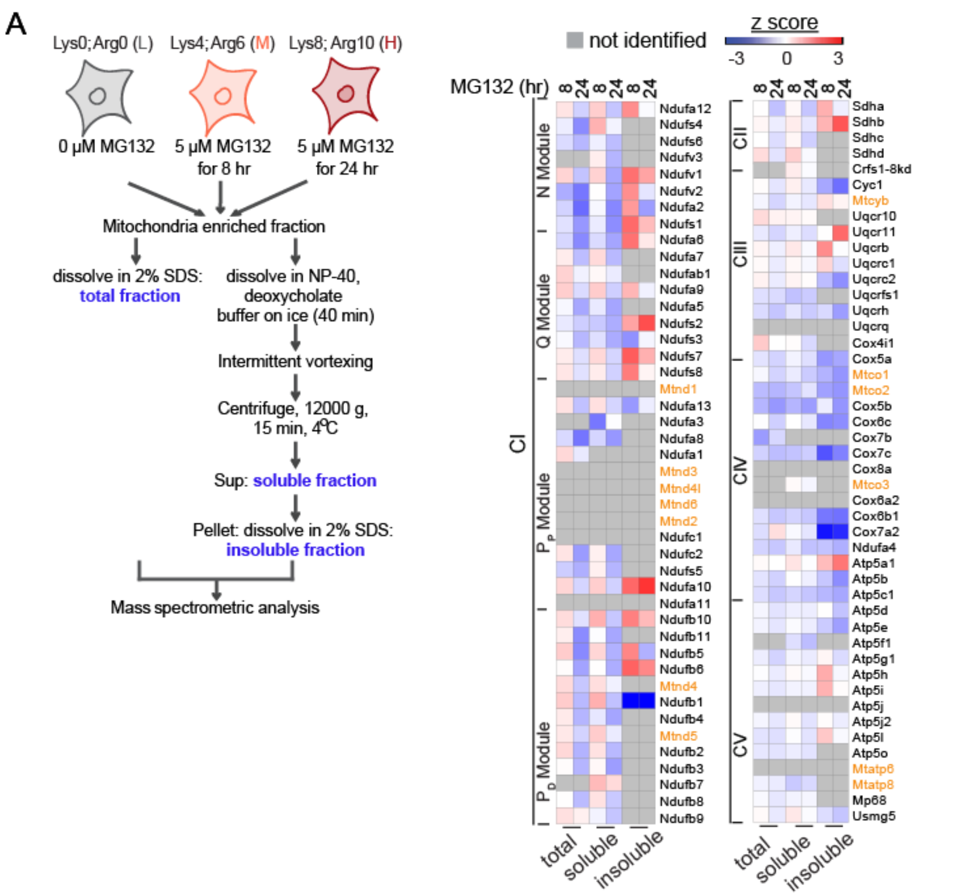


**Figure S1. RC-subunits in mitochondrial fraction during proteasome inhibition**

A) **Left:** Flowchart describing SILAC-based total, soluble and insoluble proteome profiling protocol for mitochondrial fraction of MG132 treated Neuro2a cells. **Right:** Heatmap showing distribution of z-scores calculated from SILAC fold changes for identified RC-subunits in mitochondrial fraction of MG132-treated (5µM) Neuro2a cells. Mitochondria-encoded subunits are shown in orange. n = 2.

Figure S2

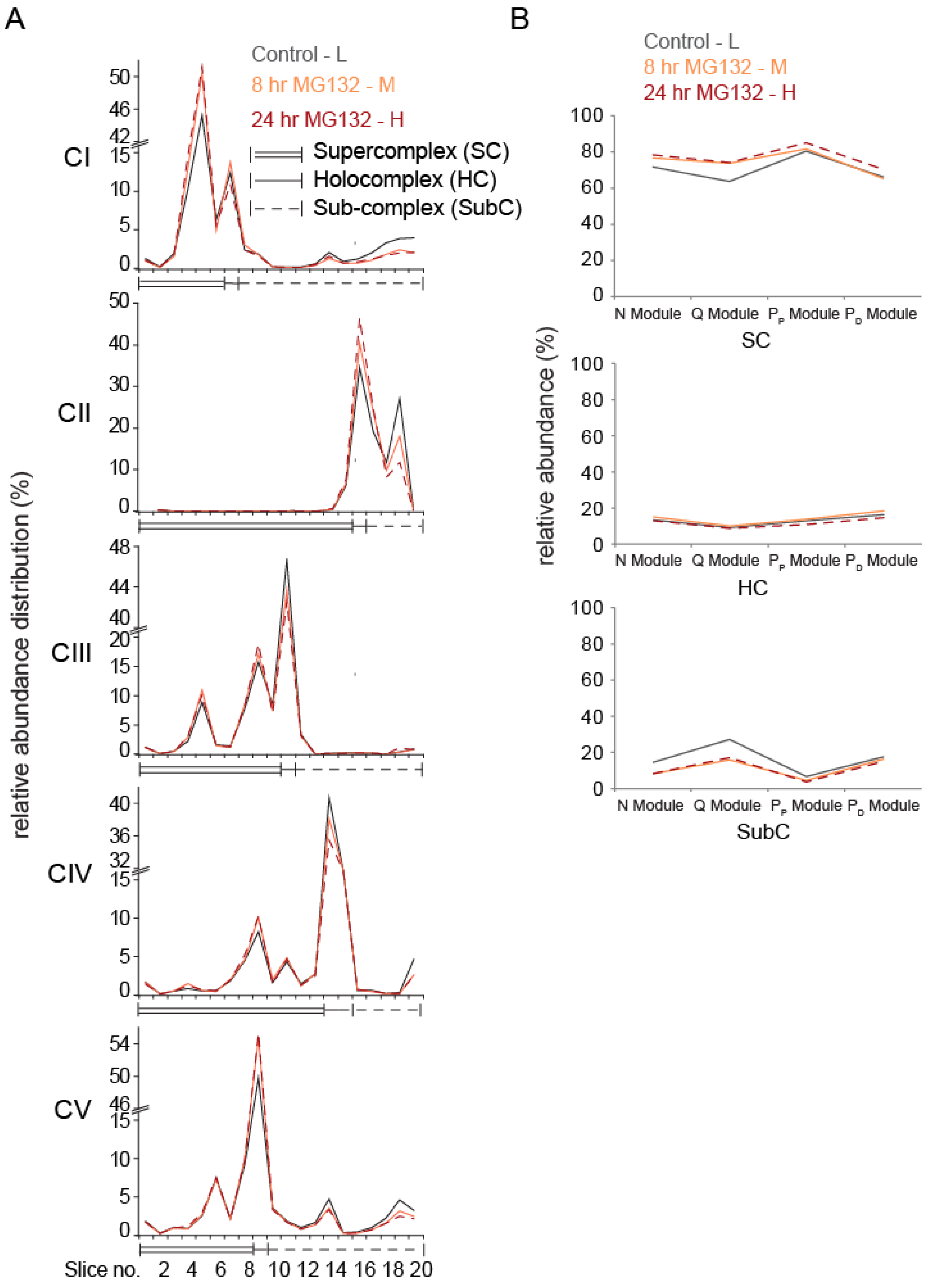


**Figure S2. Improved assembly of respiratory complexes during proteasome inhibition Neuro2a cells**

**A)** Relative abundance distribution profile of RCs across the BN-PAGE. Neuro2a cells were treated with 5 µM MG132 for 8 and 24 hr. Experimental conditions and SILAC isotopic peptides are colour coded in Figure.

**B)** Module-wise relative abundance (iBAQ) distribution of CI subunits across three sub-groups in 5 µM MG132 treated cells for 8 and 24 hr.

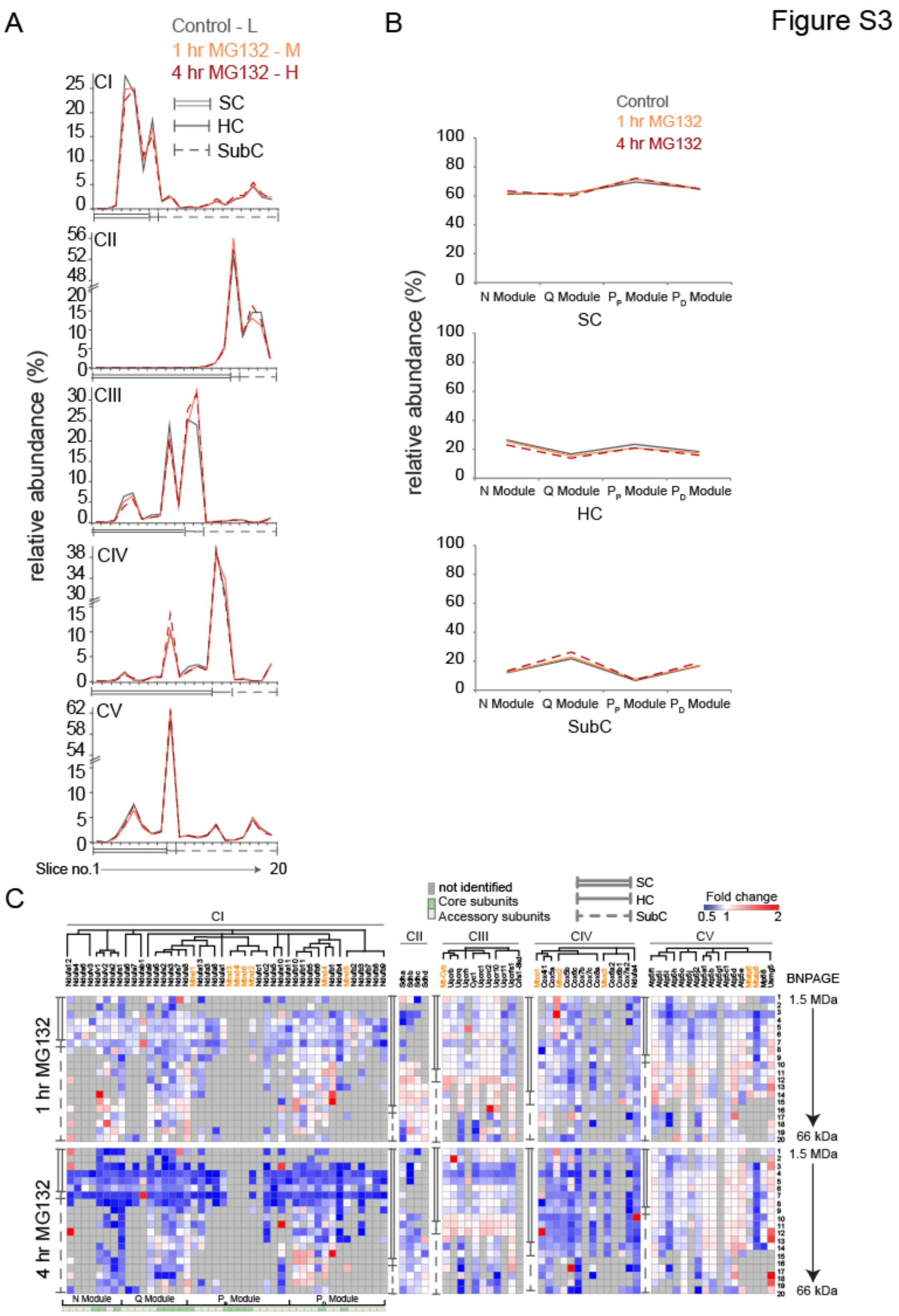


**Figure S3. iSRC at early time-points of proteasome-inhibition**

**A)** Relative abundance distribution profile of RCs across the BN-PAGE. Neuro2a cells were treated with 5 µM MG132 for 1 hr and 4 hr. Experimental conditions and SILAC isotopic peptides are colour coded in Figure. n = 2.

**B)** Module-wise relative abundance distribution of CI subunits across three sub-groups in 5 µM MG132 treated cells for 1 and 4 hr.

**C)** Heatmap showing SILAC based fold changes in abundance of RC-subunits in MG132 (5 µM) treated Neuro2A cells (1 and 4 hr) at different slices of BN PAGE. The subunits are clustered as mentioned in **Figure 2A**. Mitochondria-encoded subunits are shown in orange.

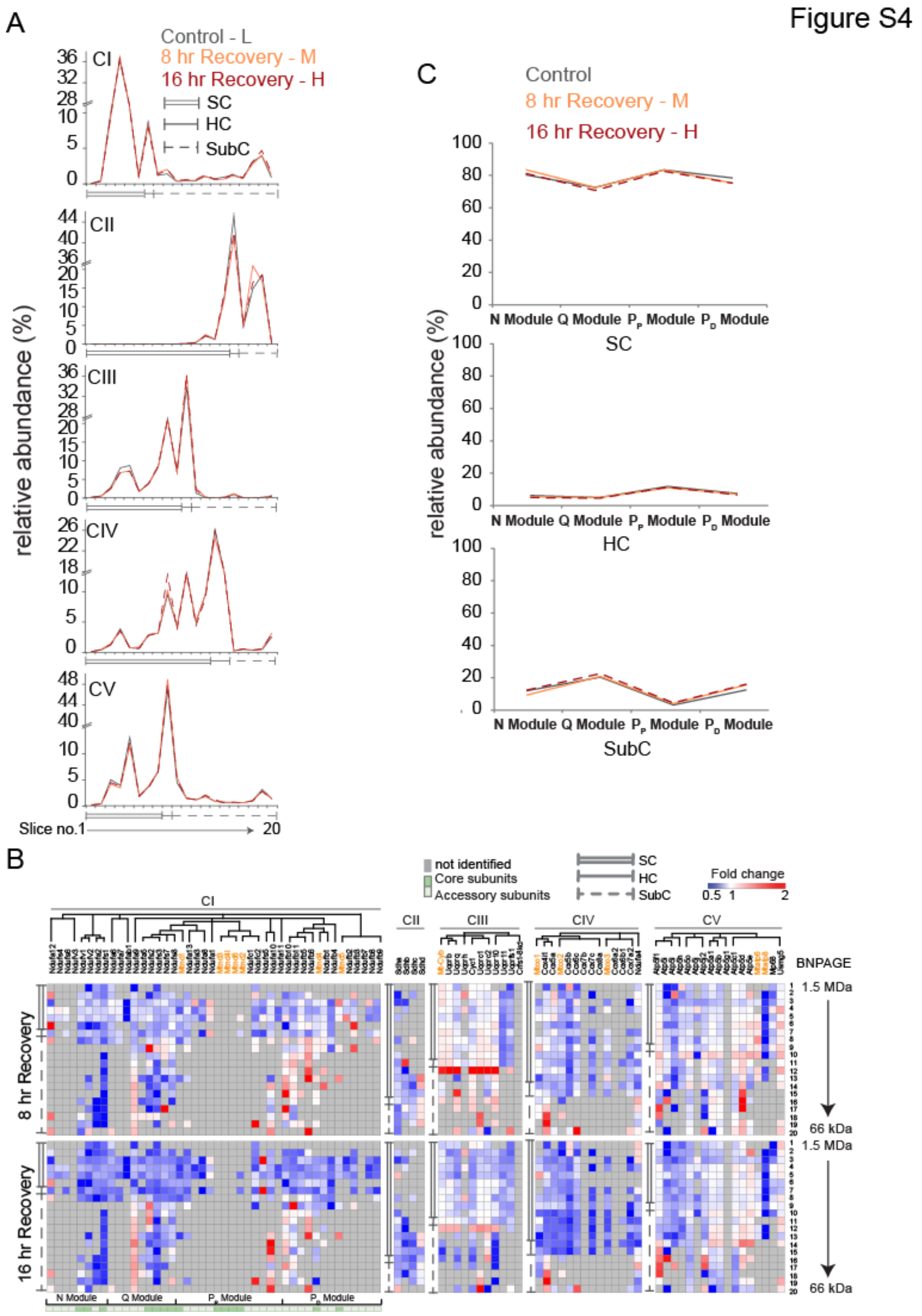

**Figure S4. Reversion of iSRC**

**A)** Relative abundance distribution profile of RCs across the BN-PAGE. Neuro2a cells were treated with 5 µM MG132 (8 hr) followed by recovery for 8 and 16 hr. Experimental conditions colour coded in Figure. n = 1.

**B)** Heatmap showing SILAC based fold changes in abundance of RC-subunits after MG132-recovery in Neuro2A cells at different slices of BN PAGE. The subunits are clustered as mentioned in **Figure 2A**. Mitochondria-encoded subunits are shown in orange.

**C)** Module-wise relative abundance distribution of CI subunits across the three sub-groups during MG132-recovery.

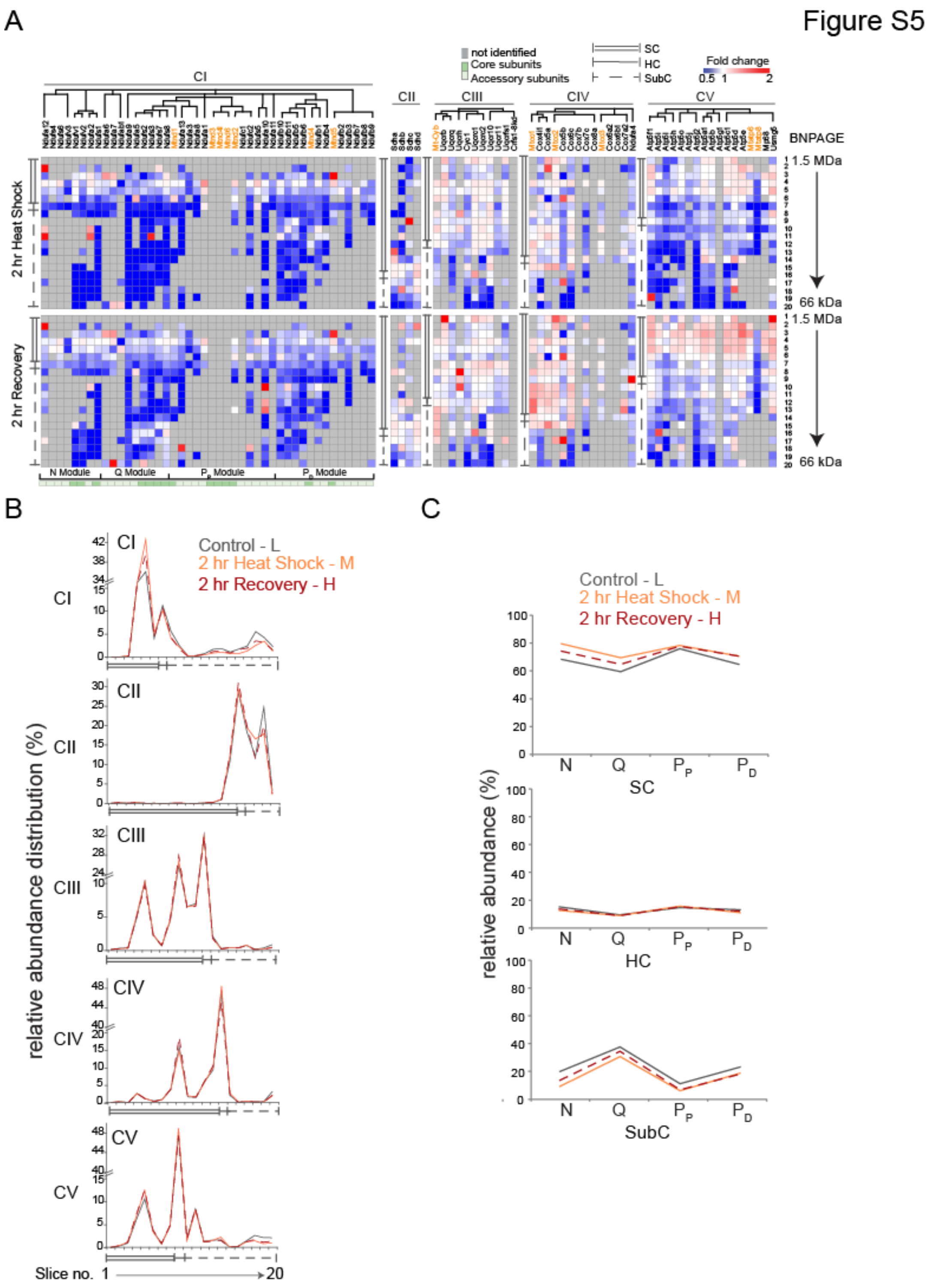


### Figure S5. iSRC in heat-stressed cells

**A)** Heatmap showing SILAC based fold changes in abundance of RC-subunits in heat-stressed Neuro2A cells (42°C for 2 hr; recovery 37°C for 2 hr) at different slices of BN

PAGE. The subunits are clustered as mentioned in **Figure 2A**. Mitochondria-encoded subunits are shown in orange. n = 1.

**B)** Relative abundance distribution profile of RCs across the BN-PAGE. Experimental conditions colour coded in Figure. n = 1.

**C)** Module-wise relative abundance distribution of CI subunits across the three sub-groups in heat stressed cells.

Figure S6

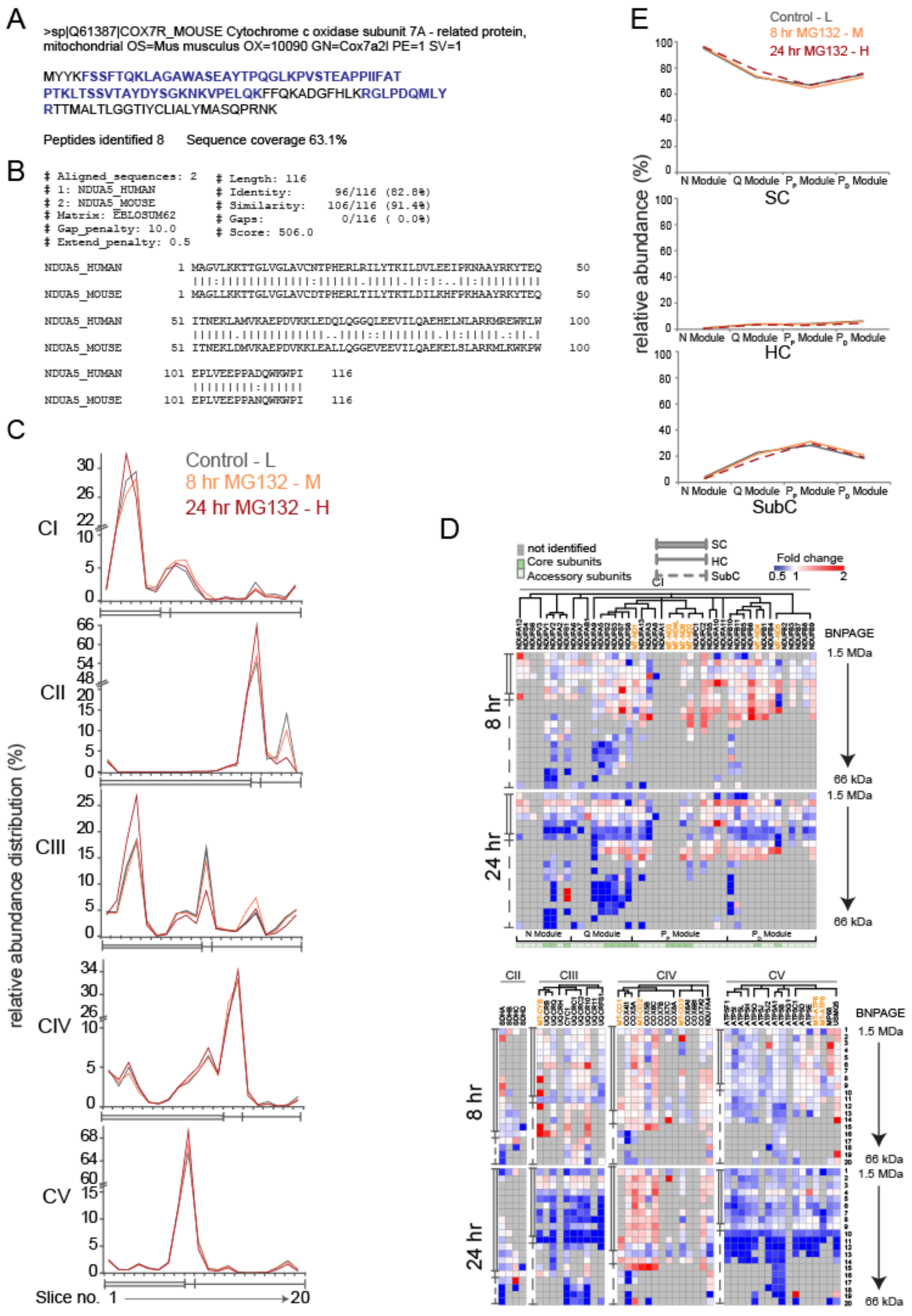

**Figure S6. iSRC in Hek293T cells**

**A)** Protein sequence of shorter isoform of Cox7a2l, supercomplex assembly factor as identified in our mass spectrometry experiment in Neuro2a cells. Identified peptide sequences are in blue and bold.

**B)** Pairwise sequence alignment of Human and Mouse Ndufa5 using EMBOSS Needle (Chojnacki et al, 2017).

**C)** Relative abundance distribution profile of RCs across the BN-PAGE. Hek293T cells were treated with 5 µM MG132 for 8 and 24 hr. Experimental conditions colour coded in Figure. n = 1.

**D)** Heatmap showing SILAC based fold changes in abundance of RC-subunits in MG132 (5 µM) treated Hek293T cells at different slices of BN PAGE. The subunits are clustered as mentioned in **Figure 2A**. Mitochondria-encoded subunits are shown in orange.

**E)** Module-wise relative abundance distribution of CI subunits across the three sub-groups in MG132-treated Hek293T cells.
